# Highly efficient cellular expression of circular mRNA enables prolonged protein expression

**DOI:** 10.1101/2023.07.11.548538

**Authors:** Mildred J. Unti, Samie R. Jaffrey

## Abstract

A major problem with mRNA therapeutics is the limited duration of protein expression due to the short half-life of mRNA. New approaches for generating highly stable circular mRNA in vitro have allowed increased duration of protein expression. However, it remains difficult to genetically encode circular mRNAs in mammalian cells, which limits the use of circular mRNA in cell-derived therapeutics. Here we describe the adaptation of the Tornado (Twister-optimized RNA for durable overexpression) system to achieve in-cell synthesis of circular mRNAs. We identify the promoter and internal ribosomal entry site (IRES) that result in high levels of protein expression in cells. We then show that these circular mRNAs can be packaged into virus-like particles (VLPs) thus enabling prolonged protein expression. Overall, these data describe a platform for synthesis of circular mRNAs and how these circular mRNAs can markedly enhance the ability of VLPs to function as a mRNA delivery tool.

## INTRODUCTION

All mRNA therapeutics have a limited duration of expression due to the relatively short half-life of mRNA in the cytoplasm. Researchers have significantly ameliorated this problem for in vitro synthesized mRNAs by synthesizing them as circular mRNAs (Wesselhoeft et al., 2018). The circular mRNAs are synthesized by ligating the 3’ and 5’ ends using enzymatic methods or permuted self-splicing introns (Obi and Chen, 2021; Puttaraju and Been, 1992; Qu et al., 2022). Circular mRNAs utilize an internal ribosome entry site (IRES) for recruiting translational machinery since they lack a 5’ cap (Chen and Sarnow, 1995). Circular RNAs are known to be highly stable (Cocquerelle et al., 1993; Jeck et al., 2013) since they cannot be degraded by exonucleases, which require 5’ or 3’ ends (Ibrahim et al., 2008). Thus, therapeutic circular mRNA can be used in place of linear mRNA to achieve prolonged expression of the encoded protein (Chen et al., 2022; Qu et al., 2022; Wesselhoeft et al., 2018).

In addition to limited duration of expression, another major challenge of mRNA therapeutics is achieving mRNA delivery to specific cell types. mRNA therapeutics are synthesized in vitro and then packaged into liposomes or lipid nanoparticles. When administered systemically, these mRNAs are taken up primarily by cells in the liver (Pardi et al., 2015). Since many applications require mRNA delivery to tissues besides the liver, an important objective is to devise strategies for the cell-type specific delivery of therapeutic mRNA beyond the liver.

One emerging approach for delivering mRNAs to specific cell types is to package mRNA into virus-like particles (VLPs). VLPs comprise the major structural proteins of a virus needed to assemble a viral capsid, but do not package viral genomic material. Instead of genomic material, VLPs can be designed to package and deliver specific mRNAs (Lu et al., 2019; Prel et al., 2015). In this case, the mRNAs are not synthesized in vitro. Instead, mRNAs are expressed in mammalian cells and directed to enter VLPs during assembly. These VLPs are produced with a nucleocapsid protein that has a MS2 coat protein (MCP) fused to the N-terminus. The nucleocapsid protein then recruits MS2 hairpin-containing mRNAs into the VLP. mRNA-containing VLPs have been used to express Cas9, Cre, and nano-luciferase (nLuc) mRNA (Lu et al., 2019; Prel et al., 2015; Segel et al., 2021).

VLPs have several advantages compared to more conventional liposome and lipid nanoparticle delivery strategies. The major advantage is that the VLPs can be “pseudotyped,” which is a process where the surface proteins are replaced with those of another virus, to modify the VLP tropism (Cronin et al., 2005; Hamilton et al., 2021; Naldini et al., 1996). An additional advantage is that VLPs deliver mRNA into the cytosol (Stein et al., 1987), rather than endosomes. When mRNA is delivered using a lipid nanoparticle, only a small amount of mRNA “escapes” from the endosome into the cytoplasm (Maugeri et al., 2019). Because VLPs deliver mRNA into the cytosol, relatively small amounts of mRNA can be used to achieve mRNA expression in target cells.

However, VLPs are limited since current methods only allow them to contain linear mRNAs. VLPs would become a more useful technology if the duration of expression of the therapeutic protein can be extended by delivering circular mRNA rather than linear mRNA. In order to achieve this, circular mRNAs need to be generated in cells and then packaged into VLPs. Previous work has found that the standard method for creating large circular RNAs, termed the backsplicing system (Liang and Wilusz, 2014), generates linear RNA species and would therefore be an inefficient method for producing VLPs that contain a circular mRNA (Ho-Xuan et al., 2020; Jiang et al., 2021).

Here we describe an improvement of VLP technology by packaging circular mRNAs rather than linear mRNAs. To do this, we developed a novel method to express circular mRNAs with high efficiency in mammalian cells. This system utilizes the Tornado (Twister-optimized RNA for durable overexpression) circular RNA expression system which has previously been used to generate high levels of small circular RNA (Litke and Jaffrey, 2019). We found that this approach produced markedly more circular mRNA in mammalian cells compared to the amount of circular mRNA produced by the backsplicing system. In a series of experiments, we then tested different promoters and IRESs to identify a combination that produced high protein output in mammalian cells—which we refer to as the “Tornado translation” system. We show that the Tornado translation system can be used to produce VLPs that contain circular mRNA, resulting in VLPs that exhibit markedly prolonged protein synthesis compared to VLPs that contain linear mRNAs. Overall, these experiments provide a new approach for delivery of circular mRNAs into target cells using VLPs.

## RESULTS

### Design of a reporter for circular mRNA-specific translation

In order to generate VLPs that contain circular mRNAs, we first needed to determine whether the Tornado translation system could be used to express circular mRNAs. To do this, we wanted to develop a system where formation of a circular mRNA produces a signal, but the precursor linear mRNA does not produce a signal. For this purpose, we used the split nano-luciferase (nLuc) system, also called the NanoLuc Binary Technology (NanoBiT), that is composed of the Large BiT (LgBiT) and Small BiT (SmBiT) components (Dixon et al., 2016). The components have low affinity for each other and so they can only produce luminescence when they are artificially brought together, for example, by two proteins or by a protein linker (**Figure 1A**).

**Figure 1.**
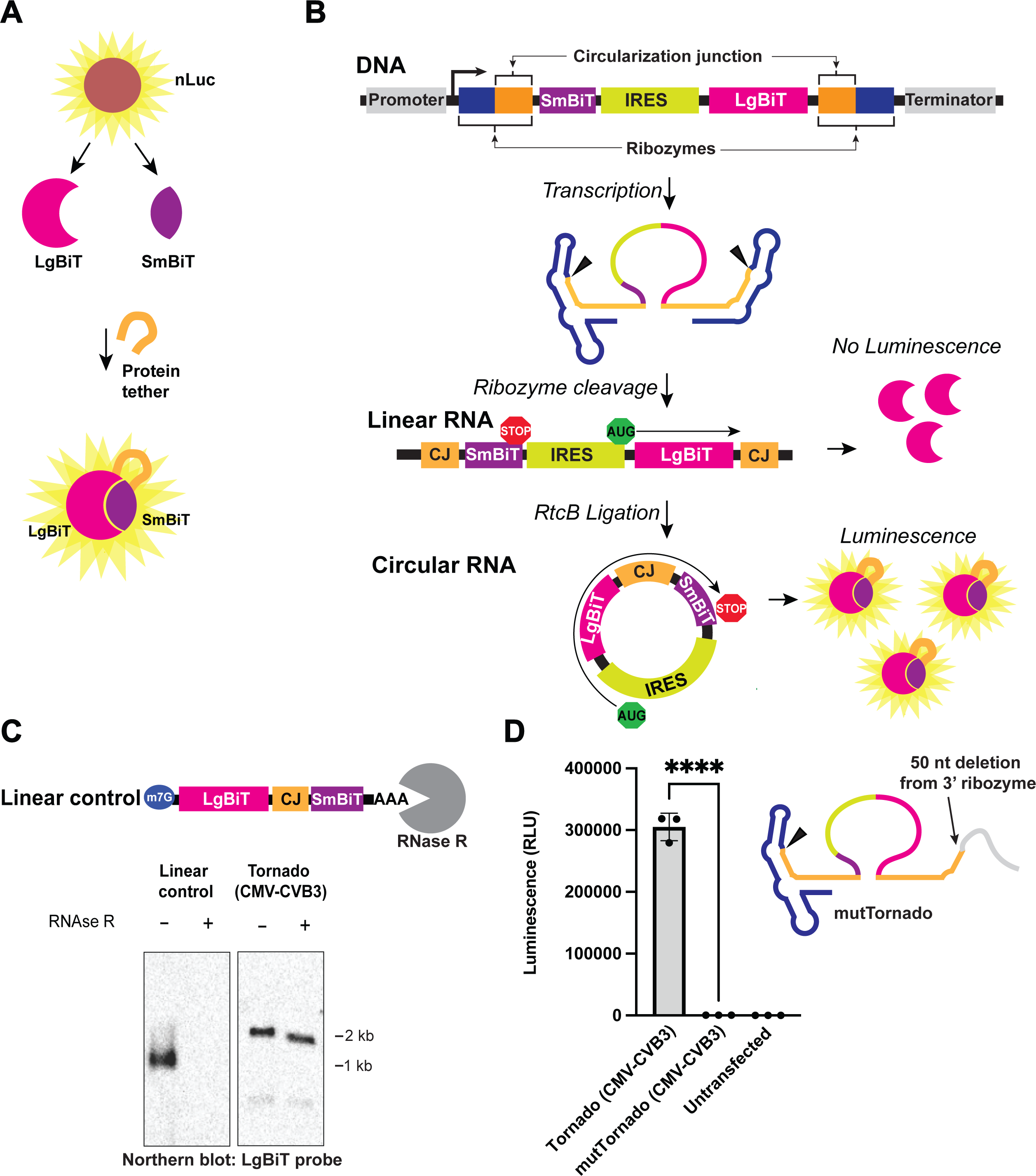
Design of a reporter for circular mRNA-specific translation. (A) Design of the split nLuc system. The LgBiT and SmBiT can only produce luminescence when they are brought together by a protein tether. (B) Construct design of the Tornado translation system using a split nLuc ORF. If the Tornado translation mRNA is in linear form, the IRES drives translation of only the LgBiT, which does not produce luminescence. If the Tornado translation mRNA is in circular form, the IRES drives translation of the LgBiT, followed by a peptide tether corresponding to the Tornado circularization junction, and then SmBiT. LgBiT tethered to the SmBiT produces luminescence. CJ = circularization junction. (C) The Tornado translation system produces a circular split nLuc mRNA. In this experiment, the Tornado translation system is expressed using a Pol II (CMV) promoter and uses a CVB3 IRES to drive translation (termed “Tornado CMV-CVB3”). Shown is a schematic of the linear construct (Linear control) used as a control for RNase R treatment. The linear RNA contains the same split nLuc ORF as the circular mRNA generated by the Tornado translation system. HEK293T cells were transfected with plasmids expressing the Tornado split nLuc mRNA (Tornado CMV-CVB3). Control cells were transfected with a plasmid expressing the linear split nLuc mRNA (Linear control). RNA was treated with vehicle or RNase R to test whether the RNA is circular. The Tornado translation system produces a circular split nLuc mRNA. Full blot image is shown in (**Figure S8**). CJ = circularization junction. (D) Luminescence is dependent on circularization of the split nLuc. We quantified luminescence from HEK293T cells transfected with a plasmid expressing the Tornado translation system with a split nLuc mRNA (Tornado (CMV-CVB3)) and a similar transcript in which the 3’ Tornado ribozyme is mutated (termed “mutTornado (CMV-CVB3)”). The mutTornado construct contains a 50 nt deletion in the 3’ ribozyme that prevents cleavage. Luminescence is only produced when both ribozymes can cleave and produce the 5’OH and 2’,3’-cyclic phosphate required for ligation by a cellular ligase. (n = 3 biological replicates). ****p<.0001, ***p<.001, **p<.01, *p<.05, n.s. p>.05. Data are presented as mean values +/− one SD.

To distinguish between translation of a circular mRNA and translation of the linear precursor, we designed mRNAs that would express SmBiT and LgBiT connected via a linker only if the RNA was circularized (**Figure 1B**). We incorporated this design into the Tornado system, which involves synthesis of a linear RNA containing ribozymes at the 5’ and 3’ ends of the transcript. After the ribozymes undergo autocatalytic cleavage, the 5’ and 3’ ends are ligated by RtcB, an endogenous RNA ligase. This ligation produces the circular RNA. The linear mRNA precursor was designed to contain three sequential components: (1) SmBiT followed by a stop codon; (2), an IRES; and lastly (3) a start codon followed by the LgBiT sequence. In this order, the linear RNA would only translate the LgBiT protein, which is unable to catalyze luminescence. However, if the mRNA is circularized, the SmBiT is now placed in a position immediately after the LgBiT. The linker between the LgBiT and the SmBiT is the circularization junction and is designed to keep LgBiT and SmBiT in frame by not having stop codons. In this way, translation can proceed from LgBiT through the circularization junction and then to SmBiT, which is terminated by a stop codon (**Figure 1B**). Luminescence will therefore only occur when the mRNA is circularized and can be translated into a protein where the circularization junction tethers the SmBiT to the LgBiT.

We next asked if this design indeed generates circular mRNA. To test this, we cloned a plasmid that expressed the Tornado translation system containing a split nLuc mRNA using the RNA polymerase II cytomegalovirus (CMV) promoter. To drive translation of the split nLuc, we included the Coxsackievirus B3 (CVB3) IRES, which has previously been used to drive translation of in vitro-transcribed circular mRNA (Wesselhoeft et al., 2018). This construct is referred to as the CMV-CVB3 Tornado translation system. We performed a northern blot using probes against the LgBiT on RNA from HEK293T cells transfected with the plasmid expressing the CMV-CVB3 Tornado translation system. The blot revealed that the Tornado translation system produced a single band (**Figure 1C**). To determine if this was the precursor linear RNA or the circular RNA, we used RNase R, which preferentially degrades linear RNA (Abe et al., 2022). We found that this band was resistant to RNase R, while a control linear RNA encoding the split nLuc was largely degraded by RNase R treatment (**Figure 1C**). These data are consistent with the premise that the Tornado translation system produces a predominantly circular mRNA. Notably, we did not detect a linear precursor RNA, similar to previous experiments with small circular RNA aptamers (Litke and Jaffrey, 2019). Since the linear mRNA precursor was not detected, the linear mRNA must have been efficiently converted to circular RNA or was degraded.

We wanted to further confirm that the luminescence derives from the circular RNA. To test this, we deleted 50 nt of the 3’ ribozyme sequence, thereby preventing the formation of the 2’,3’- cyclic phosphate at the 3’ end of the RNA, which is needed for RtcB-mediated circularization. This construct is referred to as “mutTornado”. We then compared the luciferase signal from cells transfected with the CMV-CVB3 Tornado and mutTornado translation systems. We collected samples three days after transfection to ensure the RNA had reached peak expression levels (Litke and Jaffrey, 2019) (**Figure S1A, B**). Cells transfected with the mutTornado plasmid produced no luminescence (**Figure 1D**). These data confirms that the Tornado translation split nLuc mRNA design produces a signal only when the mRNA is in circular form.

Since the Tornado system was originally designed to circularize small RNAs, we next wanted to ask whether lengthening the circularization junction would increase circular mRNA production. Circularization of the RNA is facilitated by the “circularization junction” composed of the 5’ and 3’ ends of the RNA after ribozyme cleavage. These junctional sequences hybridize to each other in order to place the 5’ and 3’ ends in proximity for RtcB-mediated ligation (**Figure S2A**). To determine whether lengthening the junctional sequence could increase circularization, we increased the length of the base paired region from 18 base pairs, which is the length in the original Tornado construct, to 26 and 49 base pairs. We measured circularization by the level of luminescence (**Figure S2A**). We found that the 26 or 49 base pair circularization junctions did not provide a statistically significant increase in luminescence compared to the 18 base pair circularization junctions (**Figure S2B**). Although not statistically significantly, the 26 base pair stem produced the most luminescence and thus was used for subsequent experiments.

### The Tornado translation system produces more circular mRNA than the backsplicing system

We next wanted to compare the amount of circular mRNA generated by the Tornado system to the that generated by the current system that is used for expressing circular mRNA in cells, i.e., the backsplicing system. The backsplicing system uses an exon comprising a gene of interest and intronic sequences from *ZKSCAN1*, a gene which normally produces a circular RNA through an endogenous backsplicing event (**Figure 2A**) (Liang and Wilusz, 2014). To directly compare the Tornado translation system to the backsplicing system, we cloned the same split nLuc open reading frame (ORF) and IRES (CVB3) from the Tornado translation system into the most used plasmid backbone for implementing the backsplicing system (Addgene #60648) (Liang and Wilusz, 2014). Surprisingly, the Tornado translation system produced ∼220-fold more luminescence than the backsplicing system (**Figure 2B**). Thus, the Tornado translation system produces markedly more circular mRNA, and therefore protein, than the backsplicing system.

**Figure 2.**
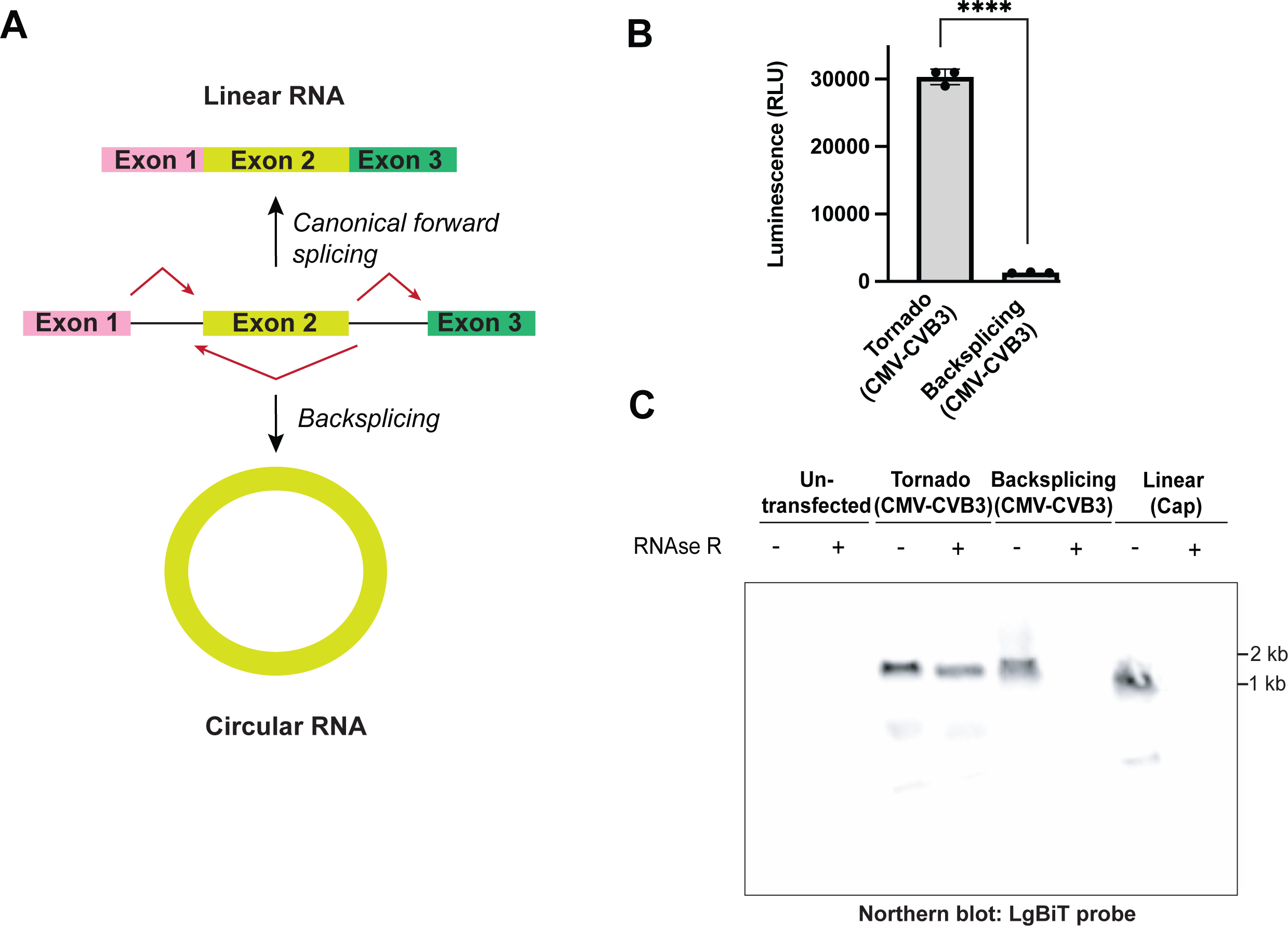
The Tornado translation system expresses more circular mRNA than the backsplicing system. (A) Schematic of the backsplicing reaction. Intron homology drives a backsplicing reaction that results in the formation of a circular RNA. (B) The Tornado translation system expresses more protein than the backsplicing system. We quantified luminescence from HEK293T cells transfected with plasmids expressing the Tornado translation (Tornado (CMV-CVB3)) and backsplicing (Backsplicing (CMV-CVB3)) systems expressing split nLuc mRNA. The Tornado translation system produced ∼220-fold more luminescence than the backsplicing system. RLU = Relative Luminescence units. Data are presented as mean values +/− one SD (n = 3 biological replicates). Significance was calculated using unpaired two-tailed student’s *t*-test. ****p<.0001, ***p<.001, **p<.01, *p<.05, n.s. p>.05. (C) The backsplicing system produces a primarily linear mRNA. HEK293T cells were transfected with plasmids expressing the Tornado translation (Tornado (CMV-CVB3)) and backsplicing (Backsplicing (CMV-CVB3)) systems. RNA was treated with vehicle or RNase R to test whether the RNA is circular. The Tornado translation system produces an mRNA that is primarily in a circular form. The backsplicing system produces an mRNA that is primarily in a linear form. Ethidium bromide-stained blot is shown in (**Figure S3B**).

Since the luminescence was much higher with the Tornado translation system, we sought to validate this result by measuring the amount of circular mRNA using northern blotting. One possibility was that the backsplicing system is producing similar levels of circular RNA as the Tornado translation system, but the two amino acid insertion into the circularization junction that was required for cloning into the backsplicing system prevented the split nLuc from folding properly (see **Supplementary information** for exact sequences used). To rule out the possibility that this two-amino acid insertion was responsible for the decreased protein signal, we measured the RNA levels generated by both systems. To do this, we performed a northern blot using probes against the LgBiT on RNA from HEK293T cells transfected with either the backsplicing or Tornado translation systems. We treated the RNA with RNase R or vehicle to identify the circular products. The Tornado translation system expressed a predominantly circular product as evidenced by the presence of an RNase R resistant band, whereas the backsplicing system created a predominantly linear band as evidenced by the disappearance of the band after RNAse R treatment (**Figure 2C**). These data suggested that the major product of the backsplicing system is a linear RNA.

As an additional control to confirm that the backsplicing system produces a predominantly linear RNA, we used the endogenous *ZKSCAN1* sequence. In these experiments we measured the circular RNA production using the *ZKSCAN1* exon2/3 insert cloned into the Tornado system and the most used plasmid backbone for implementing the backsplicing system (Liang and Wilusz, 2014). To measure circular RNA production, we performed a northern blot using probes against the *ZKSCAN1* exons 2/3 on RNA from HEK293T cells transfected with either the backsplicing or Tornado translation systems. We treated the RNA with RNase R or vehicle to identify the circular products. As expected, we found that the Tornado translation system created a single predominant band, which was a circular RNA based on its resistance to RNAse R. However, the backsplicing system created a predominantly linear band as evidenced by the disappearance of the band after RNAse R treatment (**Figure S3A**). This result indicates that the Tornado translation system produces markedly more circular RNA compared to the backsplicing system, and that this effect is not insert-specific.

Notably, two studies recently reported that the backsplicing system expresses several linear transcripts (Ho-Xuan et al., 2020; Jiang et al., 2021). Thus, the backsplicing system produces most of its RNA in a linear form while the Tornado system produces predominantly circular mRNA. These data suggest that the Tornado translation system is the superior method for cell-based synthesis of circular mRNAs for VLPs or other applications.

### Selection of the Pol II-compatible IRES

We next wanted to identify a promoter and IRES that would produce high-level protein expression from a Tornado-expressed circular mRNA. A commonly used IRES is the encephalomyocarditis virus (EMCV) IRES (Jang and Wimmer, 1990). However, a recent study showed that the CVB3 IRES produces more protein than the EMCV IRES when used in the context of an in vitro-transcribed circular mRNA (Wesselhoeft et al., 2018). A more recent study found that the related IRES derived from the human rhinovirus B3 (HRV-B3) provides substantially more protein production than the CVB3 IRES (Chen et al., 2022). We therefore compared these IRESs in the Tornado translation system. In these experiments, only the IRES was changed. Each construct was expressed from a Pol II (CMV) promoter, producing a transcript encoding the split nLuc. We found that the CVB3 IRES produces more protein than the EMCV IRES and slightly more protein than the HRV-B3 IRES (**Figure 3A**, **Figure S4**). Thus, the CVB3 IRES produces the highest level of protein expression from a Pol II-driven Tornado translation system.

**Figure 3.**
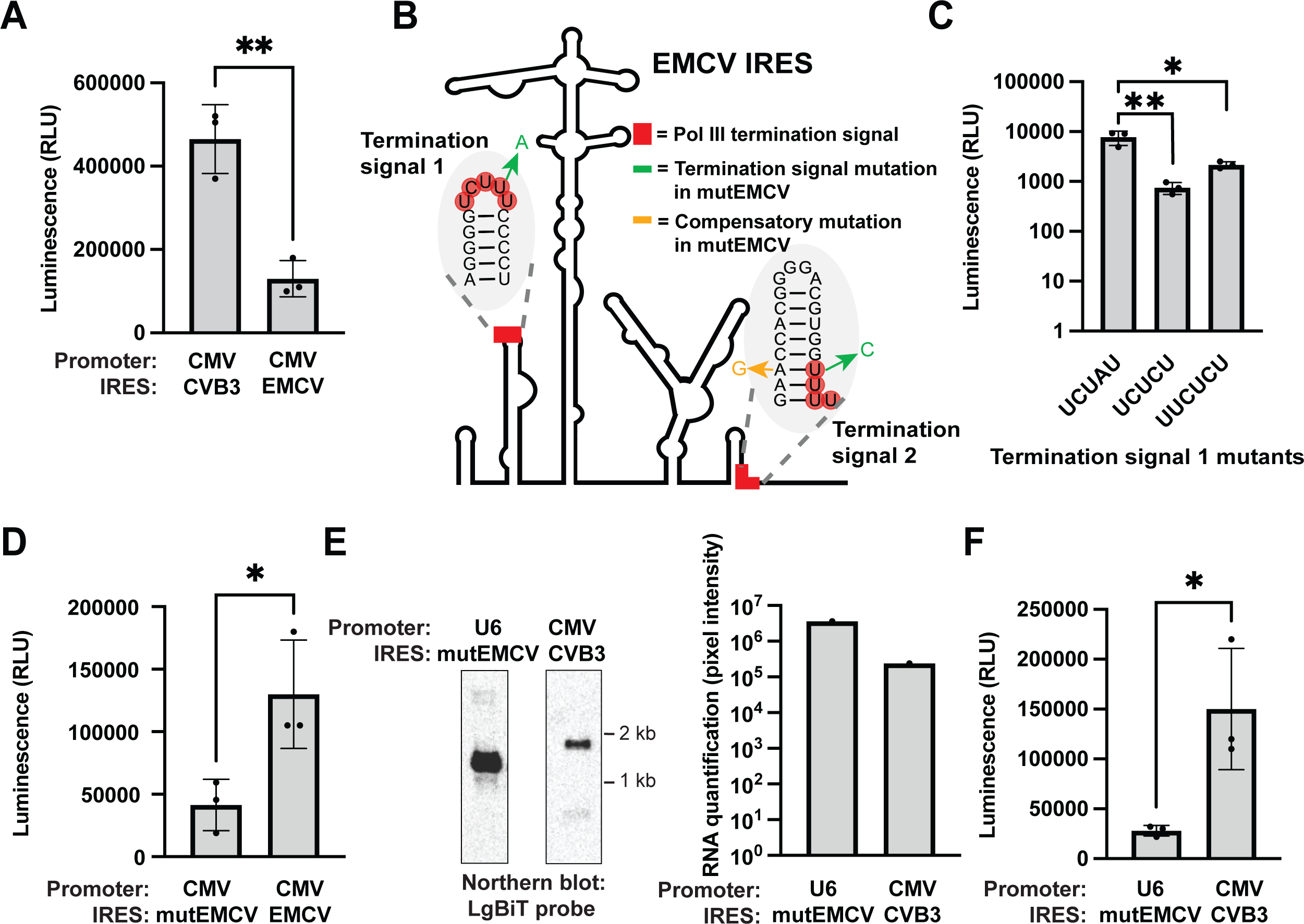
The Tornado translation system produces the most protein using a CMV-CVB3 promoter and IRES combination. (A) The CVB3 IRES expresses more protein than the EMCV IRES. We quantified luminescence from HEK293T cells transfected with plasmids expressing the CVB3 or EMCV IRES. Both constructs were expressed using a Pol II-driven (CMV) Tornado translation system with the split nLuc ORF. The CVB3 IRES produced ∼3.5-fold more luminescence than the EMCV IRES. RLU = Relative Luminescence units. Data are presented as mean values +/− one SD (n = 3 biological replicates). Significance was calculated using unpaired two-tailed student’s *t*-test. ****p<.0001, ***p<.001, **p<.01, *p<.05, n.s. p>.05. (B) Mutations required to create Pol III-compatible EMCV IRES. Pol III termination signals are shown in red. Mutations that render the IRES compatible with a Pol III promoter are shown in green, and the compensatory mutations to conserve the function of the IRES are shown in yellow. The mutant EMCV (termed “mutEMCV”) contains the mutations shown in green and yellow for termination signal 1 and 2. (C) The UCUAU termination signal 1 EMCV IRES mutant expresses more protein than the UCUCU and UUCUCU mutants. We quantified luminescence from HEK293T cells transfected with plasmids expressing EMCV IRES mutants. Both constructs were expressed using a Pol II-driven (CMV) Tornado translation system with the split nLuc ORF. The UCUAU mutant produced ∼10-fold more luminescence than the UCUCU mutant and ∼3.5-fold more luminescence than the UUCUCU mutant. The UCUAU mutation was used in the mutant EMCV (mutEMCV) IRES along with the termination signal 2 mutation. RLU = Relative Luminescence units. Data are presented as mean values +/− one SD (n = 3 biological replicates). Significance was calculated using unpaired two-tailed student’s *t*-test. ****p<.0001, ***p<.001, **p<.01, *p<.05, n.s. p>.05. (D) Mutant EMCV IRES expresses less protein than the wild-type EMCV IRES. We quantified luminescence from HEK293T cells transfected with plasmids expressing the mutant EMCV (mutEMCV) and wild-type EMCV (EMCV) IRESs. Both constructs were expressed using a Pol II-driven (CMV) Tornado translation system with the split nLuc ORF. The wild-type EMCV IRES produced ∼3-fold more luminescence than the mutant EMCV IRES. RLU = Relative Luminescence units. Data are presented as mean values +/− one SD (n = 3 biological replicates). Significance was calculated using unpaired two-tailed student’s *t*-test. ****p<.0001, ***p<.001, **p<.01, *p<.05, n.s. p>.05. (E) The Pol III-driven Tornado translation system produces markedly more RNA than the Pol II-driven Tornado translation system. HEK293T cells were transfected with plasmids expressing Pol II-driven (CMV CVB3) and Pol III-driven (U6 mutEMCV) Tornado translation systems. RNA was quantified by performing a northern blot and calculating pixel intensity with the ImageLab software (**Figure S8**). Full blot image is shown in (**Figure S8**). The Pol III-driven Tornado translation system produced ∼10-fold more RNA than the Pol II-driven Tornado translation system. (F) Despite lower RNA expression, the Pol II-driven Tornado translation system produces more protein than the Pol III-driven Tornado translation system. We quantified luminescence from HEK293T cells transfected with plasmids expressing Pol II-driven (CMV CVB3) and Pol III-driven (U6 mutEMCV) Tornado translation systems with a split nLuc ORF. The Pol II-driven Tornado translation system produced ∼6-fold more luminescence than the Pol III-driven Tornado translation system. The improved performance of the Pol II-driven Tornado translation system is likely due, in part, to the improved function of the non-mutated IRES that can be used with the Pol II promoter. RLU = Relative Luminescence units. Data are presented as mean values +/− one SD (n = 3 biological replicates). Significance was calculated using unpaired two-tailed student’s *t*-test. ****p<.0001, ***p<.001, **p<.01, *p<.05, n.s. p>.05.

**Figure 4.**
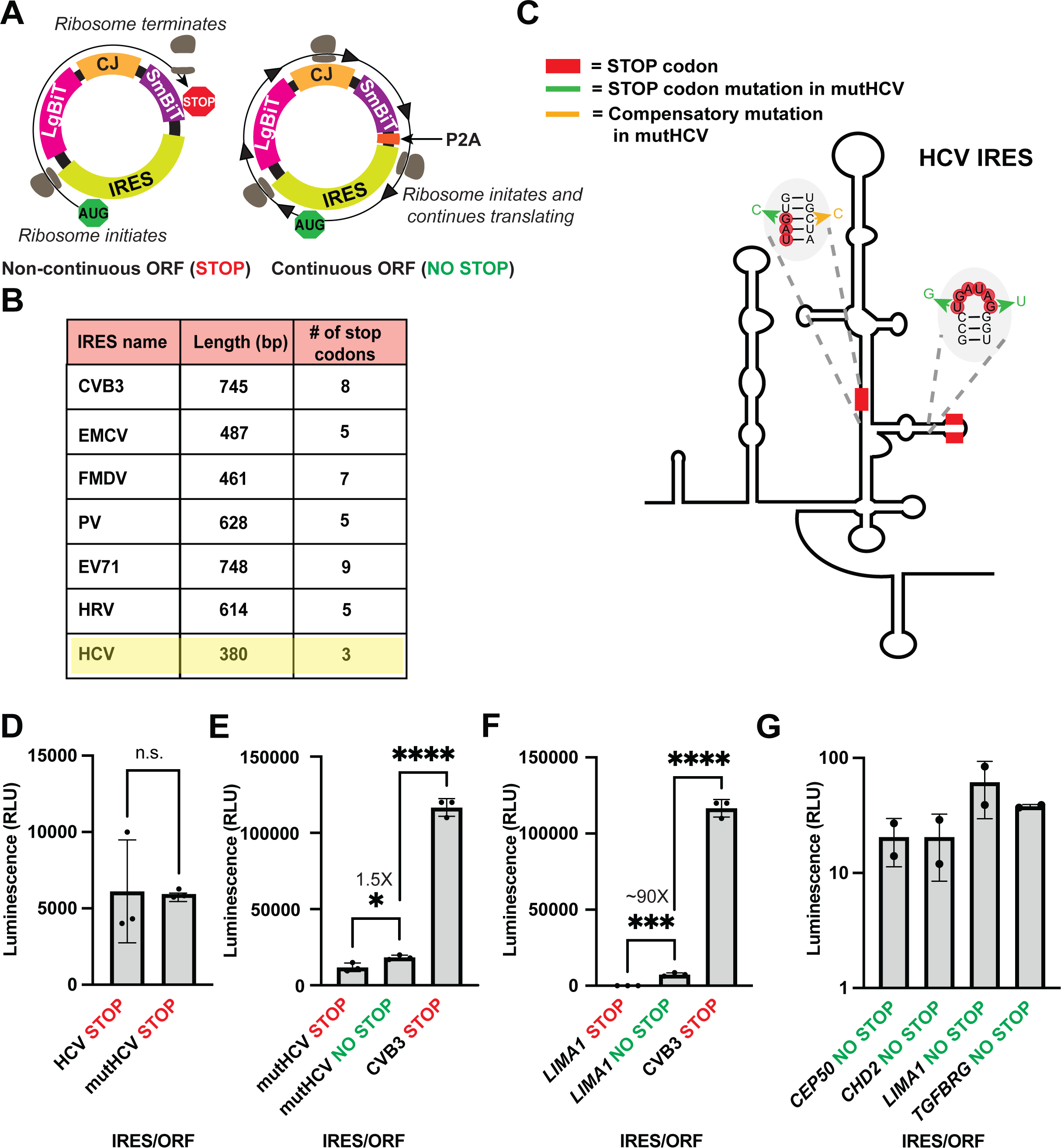
Continuous translation does not improve the protein output from the Tornado translation system. (A) Design of a non-continuous and continuous translation system. A continuous ORF does not contain a stop codon and the ribosome therefore continuously translates around the circular RNA. The P2A cleavage site cleaves the polypeptide into protein monomers. (B) Viral IRESs contain several stop codons in all frames. The HCV IRES is the IRES with the shortest length and has the fewest stop codons in any frame, and was therefore selected for mutagenesis for assessing continuous translation. (C) Mutations required to make the HCV IRES compatible with continuous translation. The sequences of the three stop codons are shown in red. Mutations that render the IRES compatible with continuous translation are shown in green, and the compensatory mutations to conserve the function of the IRES are shown in yellow. The mutant HCV (termed “mutHCV”) contains the mutations shown in green and yellow. (D) The mutHCV IRES retains translational activity. We quantified luminescence from HEK293T cells transfected with plasmids expressing the mutHCV non-continuous translation system (mutHCV STOP) and the wild-type HCV (termed “wtHCV”) non-continuous translation system (wtHCV STOP). Both constructs were expressed using a Pol II-driven (CMV) Tornado translation system with the split nLuc ORF. The mutHCV produced similar luminescence as the wtHCV. RLU = Relative Luminescence units. Data are presented as mean values +/− one SD (n = 3 biological replicates). Significance was calculated using unpaired two-tailed student’s *t*-test. ****p<.0001, ***p<.001, **p<.01, *p<.05, n.s. p>.05. (E) The mutHCV IRES does not efficiently drive continuous translation and does not produce more protein than the CVB3 non-continuous translation system. We quantified luminescence from HEK293T cells transfected with plasmids expressing the mutHCV non-continuous translation system (mutHCV STOP), the mutHCV continuous translation system (mutHCV NO STOP), and the CVB3 non-continuous translation system (CVB3 STOP). All constructs were expressed using a Pol II-driven (CMV) Tornado translation system with the split nLuc ORF. The continuous mutHCV translation system produced ∼1.5-fold more luminescence than the non-continuous mutHCV translation system. The CVB3 non-continuous translation system produced ∼10-fold more luminescence than the mutHCV continuous translation system. RLU = Relative Luminescence units. Data are presented as mean values +/− one SD (n = 3 biological replicates). Significance was calculated using unpaired two-tailed student’s *t*-test. ****p<.0001, ***p<.001, **p<.01, *p<.05, n.s. p>.05. (F) The *LIMA1* IRES can drive continuous translation but even the continuous *LIMA1* system does not produce more protein than the CVB3 non-continuous translation system. We quantified luminescence from HEK293T cells transfected with plasmids expressing the *LIMA1* non-continuous translation system (*LIMA1* STOP), the *LIMA1* continuous translation system (*LIMA1* NO STOP), and the CVB3 non-continuous translation system (CVB3 STOP). All constructs were expressed using a Pol II-driven (CMV) Tornado translation system with the split nLuc ORF. The *LIMA1* continuous translation system produced ∼90-fold more luminescence than the *LIMA1* non-continuous translation system. However, the CVB3 non-continuous translation system produced ∼15-fold more luminescence than the *LIMA1* continuous translation system. RLU = Relative Luminescence units. Data are presented as mean values +/− one SD (n = 3 biological replicates). Significance was calculated using unpaired two-tailed student’s *t*-test. ****p<.0001, ***p<.001, **p<.01, *p<.05, n.s. p>.05. (G) Alternative IRESs elements from (Chen et al., 2021) do not produce more protein than the *LIMA1* IRES. We quantified luminescence from HEK293T cells transfected with plasmids expressing the putative IRESs from the screen by (Chen et al., 2021). All constructs were expressed using a Pol II-driven (CMV) Tornado translation system with the continuous split nLuc ORF. None of the IRES elements from (Chen et al., 2021) that we tested produced more protein than the *LIMA1* IRES. RLU = Relative Luminescence units. Data are presented as mean values +/− one SD (n = 2 technical replicates). Significance was calculated using unpaired two-tailed student’s *t*-test. ****p<.0001, ***p<.001, **p<.01, *p<.05, n.s. p>.05.

### Engineering a Pol III-compatible IRES

In the experiments above, we expressed the Tornado translation system using a Pol II CMV promoter. The original Tornado expression system for small circular RNAs used a U6 Pol III promoter (Litke and Jaffrey, 2019), which is often used for expressing small RNAs (Mäkinen et al., 2006). RNA Pol III is considered a highly efficient expression system due to polymerase re-entry into the promoter after transcription termination (Dieci and Sentenac, 1996). The Tornado translation system could benefit by using a Pol III promoter because Pol III promoters express higher levels of RNA than a Pol II promoter (Dieci and Sentenac, 1996). We therefore asked whether large circular RNAs can be produced using a Pol III promoter.

A problem with using a Pol III promoter is that the commonly used IRESs such as CVB3 and EMCV contain Pol III termination signals and are therefore not expected to be transcribed by Pol III. The Pol III termination signal sequence comprises UUUU or a closely related sequence such as UCUUU or UUUAU (Orioli et al., 2011). This termination signal is quite simple and therefore occurs relatively frequently. The EMCV, CVB3, and HRV-B3 IRESs have two, three, and four Pol III termination signals respectively (**Figure 3B**, **Figure S5**). Importantly, the CVB3 and HRV-B3 IRESs contain a highly conserved U-stretch that is responsible for binding directly to 18S rRNA (Bailey Jennifer and Tapprich William, 2007; Yang et al., 2003). Mutational analysis has revealed that any mutation of this sequence will lead to abrogation of IRES activity (Yang et al., 2003). It is therefore difficult to create an IRES that is compatible with Pol III due to the presence of multiple Pol III termination signals.

**Figure 5.**
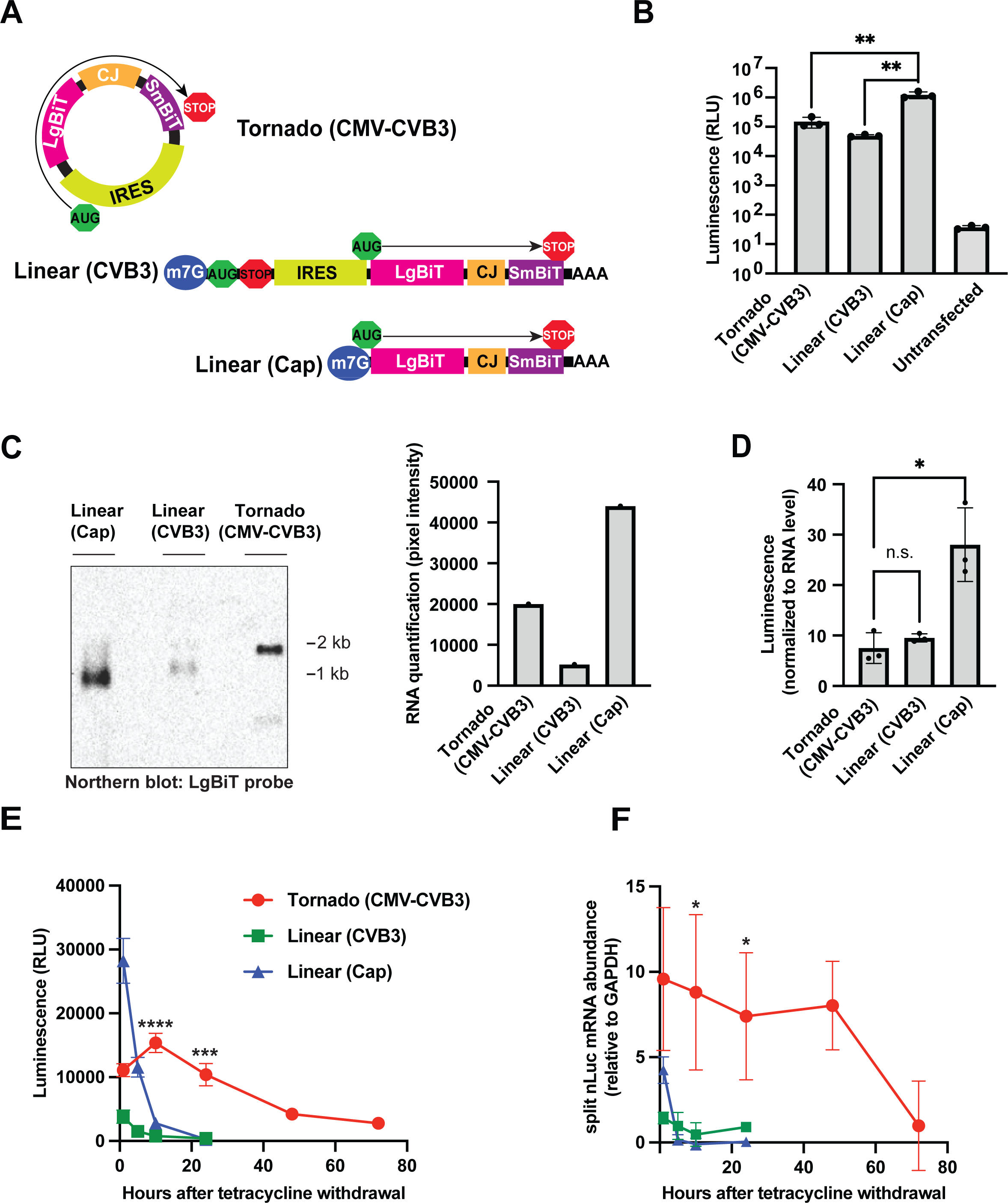
The Tornado translation system produces lower, but more persistent levels of protein production than a linear mRNA translation system. (A) Construct design of Tornado translation and linear mRNA expression systems. All three mRNAs contain the same ORF. The LgBiT is tethered to the SmBiT by the peptide that corresponds to the Tornado circularization junction. The “Linear (CVB3)” mRNA contains a start and stop codon upstream of the CVB3 IRES to ensure the translation of the split nLuc ORF is CVB3-dependent. The “Linear (Cap)” mRNA is driven by cap-dependent translation. CJ = Circularization junction. (B) The Tornado translation system produces more protein than the linear CVB3-dependent translation system but less protein than the linear cap-dependent translation system. We quantified luminescence from HEK293T cells transfected with plasmids expressing the Tornado translation system (Tornado (CMV-CVB3)), the linear cap-dependent mRNA expression system (Linear (Cap)), and the linear CVB3-dependent mRNA expression system (Linear (CVB3)). The Tornado translation system produced ∼3-fold more luminescence than the linear CVB3-dependent mRNA expression system and ∼5-fold less luminescence than the linear cap-dependent mRNA expression system. RLU = Relative Luminescence units. Data are presented as mean values +/− one SD (n = 3 biological replicates). Significance was calculated using unpaired two-tailed student’s *t*-test. ****p<.0001, ***p<.001, **p<.01, *p<.05, n.s. p>.05. (C) The CVB3 IRES leads to lower levels of RNA. HEK293T cells were transfected with plasmids expressing the Tornado translation system (Tornado (CMV-CVB3)), the linear cap-dependent mRNA expression system (Linear (Cap)), and the linear CVB3-dependent mRNA expression system (Linear (CVB3)). RNA was quantified by performing a northern blot and calculating pixel intensity with the ImageLab software (**Figure S8**). Full blot image is shown in (**Figure S8**). The linear cap-dependent mRNA expression system produced ∼3-fold more RNA than the Tornado translation system and ∼8.5-fold more RNA than the linear CVB3-dependent mRNA expression system. (D) CVB3-dependent translation produces less protein than cap-dependent translation. We normalized luminescence from (**b**) to RNA expression from (**c**). CVB3-dependent translation (Tornado (CMV-CVB3) and (Linear (CVB3)) produced ∼3-fold less luminescence than cap-dependent translation. (E, F) The Tornado translation system expresses an mRNA with a markedly longer half-life than the linear expression systems. We generated stable cells lines expressing the linear cap-dependent mRNA expression system (Linear (Cap)) (blue), the linear CVB3-dependent mRNA expression system (Linear (CVB3)) (green), and Tornado translation system (Tornado (CMV-CVB3)) (red) under a tetracycline-responsive promoter. We then pulsed these cell lines with tetracycline (1 μg/mL) for 12 hours then measured luminescence and RNA abundance periodically after replacing the media with tetracycline-free media. We quantified RNA by performing qRT-PCR with primers that amplify a 120 nt region of the LgBiT. We used *GAPDH* primers to normalize RNA abundance, then corrected for cell division (see **Method Details**). The Tornado translation system (Tornado (CMV-CVB3)) exhibits ∼10-fold longer duration of protein and mRNA expression compared to the linear mRNA expression systems. RLU = Relative Luminescence units. Data are presented as mean values +/− one SD (n = 3 biological replicates). Significance was calculated using unpaired two-tailed student’s *t*-test. ****p<.0001, ***p<.001, **p<.01, *p<.05, n.s. p>.05. CJ = circularization junction.

Because the CVB3 IRES contains an essential 18S-binding U-stretch that constitutes a Pol III termination signal, we instead focused on the EMCV IRES. We attempted to create a Pol III-compatible EMCV IRES by mutating the Pol III termination signals. The EMCV IRES contains two Pol III termination signals (**Figure 3B**). The first Pol III termination signal comprises a sequence (UCUUU) that binds polypyrimidine-tract-binding protein (PTB), which is thought to be important for IRES function by recruiting translation initiation factors (**Figure 3B**) (Kaminski and Jackson, 1998). To maintain the function of the PTB binding site but mutate the Pol III termination signal, we used alternative PTB-binding sites that do not contain a Pol III termination signal. Previous studies have shown that PTB can bind several U-rich sequences, with the most common sequence being UUCUCU (Xue et al., 2009). UUCUCU is not a Pol III termination signal (Orioli et al., 2011), so we replaced the Pol III termination element with UUCUCU. In addition, we also created an EMCV variant in which this sequence was mutated to UCUCU, which has also been described as a PTB-binding motif (Xue et al., 2009), and a UCUAU sequence, which is not a canonical PTB-binding motif (Xue et al., 2009) (**Figure 3B**). We then compared these EMCV variants by cloning them upstream of the split nLuc ORF. Surprisingly, the UCUAU mutant produced the most luminescence despite not being a canonical PTB-binding sequence (**Figure 3C**). We therefore selected the UCUAU mutation to use in the Pol III-compatible EMCV (mutEMCV) IRES.

To mutate the second Pol III termination signal we used a BLAST alignment to identify a related IRES, the Falcon picornavirus, that has a related sequence but no Pol III termination element in this region (**Figure S6A, B**). We then used the Falcon picornavirus sequence in the mutEMCV IRES in place of the Pol III termination signal in the wild-type EMCV (wtEMCV) IRES (**Figure 3B**).

We next wanted to ask whether the mutEMCV IRES had decreased translational activity compared to the wtEMCV IRES. To do this, we compared the protein output of the mutEMCV IRES to the wtEMCV IRES using a Pol II promoter, which transcribes regardless of the Pol III termination elements. We found that the wtEMCV IRES produced ∼3-fold more luminescence than mutEMCV IRES (**Figure 3D**). Thus, the mutations decreased the translational activity of the IRES, but the mutEMCV IRES still maintains translational activity.

Lastly, we wanted to compare the translational activity of the mutEMCV IRES to an IRES that naturally does not contain a Pol-III termination signal. The classical swine fever virus (CSFV) IRES is an Hepatitis C virus (HCV)-like IRES yet lacks the Pol III termination element that is present in the HCV IRES (**Figure S7A**) (Sizova Daria et al., 1998). While previous studies have found that the CSFV IRES translates ∼2-fold less protein than the EMCV IRES (Wesselhoeft et al., 2018), it does not require any mutations that decrease its function to make it compatible with a Pol III promoter. We therefore compared the mutEMCV IRES to the CSFV IRES using the Pol III-driven (U6) Tornado translation system. The mutEMCV IRES provides >3-fold more luminescence than the CSFV IRES (**Figure S7B**). This further establishes that the mutEMCV is currently the Pol III-compatible IRES with the highest translational activity.

### The Tornado translation system produces the most protein using a CMV-CVB3 promoter and IRES combination

We next asked whether protein expression is more efficient using the Pol II-driven (CMV) Tornado translation system, which relies on the wild-type IRES, or the Pol III-driven (U6) Tornado translation system, which relies on the mutated IRES.

As a first test, we examined the amount of circular mRNA produced by both systems. To do this, we performed a northern blot using probes against the LgBiT on RNA from HEK293T cells transfected with either the Pol II- or Pol III-driven Tornado translation system. As expected, a single prominent band of the same molecular weight was seen upon transfection with either construct (**Figure 3E**). However, the Pol III-driven (U6) Tornado translation system expressed over ∼10-fold more RNA than the Pol II-driven (CMV) Tornado translation system (**Figure 3E**). Overall, these results show that the Pol III-driven system generates substantially more circular mRNA than the Pol II-driven system.

We next compared the protein expression from the Pol II- and Pol III-driven Tornado translation systems. The luminescence from the Pol II-driven system was ∼6-fold higher than the Pol III-driven system (**Figure 3F**). This is in part due to the decreased translational activity of the mutEMCV IRES (**Figure 3D**). However, we would expect the ∼10-fold higher RNA levels from the Pol III system to cause higher protein expression despite the ∼3-fold decreased translational activity of the mutEMCV IRES. The lower protein output from the Pol III-driven system therefore results from a combination of reduced translation from the mutEMCV IRES and other unknown factors associated with Pol III transcription. Thus, if the goal is to express large amounts of circular RNA, the Pol III promoter should be used. However, if the goal is to express large amounts of protein from a circular mRNA, the Pol II (CMV) system with the wild-type CVB3 IRES should be used.

### Continuous translation does not improve the protein output from the Tornado translation system

We next asked if the protein production could be further increased using continuous translation. Continuous translation is when the ORF lacks a stop codon, and the ribosome continues translating past the ORF onto the IRES, then back onto the ORF (Abe et al., 2013). Continuous translation therefore requires an IRES that is in frame with the ORF and designed to not have a stop codon. In this way, the ribosome would conceivably circumambulate the circular RNA indefinitely (**Figure 4A**) (Abe et al., 2015; Costello et al., 2019). Inclusion of a viral P2A sequence, which induces ribosome skipping and therefore cleaves the polypeptide chain (Liu et al., 2017), can ensure that the protein polymer created by continuous translation is separated into functional protein monomers. Continuous translation bypasses the rate limiting step of translation, ribosome initiation (Shah et al., 2013). Therefore, continuous translation can theoretically produce more protein than a non-continuous ORF.

A problem with implementing a continuous translation system is that most IRES sequences contain many stop codons in all three reading frames that would prevent continuous translation from occurring (**Figure 4B**). Of the common viral IRES sequences, the HCV IRES contains only three stop codons in one of its reading frames, which is the fewest stop codons of any of the examined IRESs in any frame (**Figure 4B**). The first stop codon occurs in a stem whose structure, but not sequence, is conserved in related IRESs (Honda et al., 1999). Thus, we made a UAG to UAC mutation and a mutation on the complementary base to conserve the structure of the stem (**Figure 4C**). The second two stop codons are adjacent (<u>UGA UAG</u>) and contain a conserved loop sequence (GAUA) (Brown et al., 1992) (**Figure 4C**). We mutated the loop from UGAUAG to GGAUAU in order to maintain both the sequence of the conserved region and the G-U wobble base pair at the base of the loop, while removing both stop codons (**Figure 4C**). These two mutations were used to create the mutant HCV (mutHCV) IRES that could be used for continuous translation since the mutations create a continuous ORF within the IRES.

We next wanted to ask whether the mutHCV IRES had decreased translational activity compared to the wild-type HCV (wtHCV) IRES. To test this, we cloned the mutHCV and the wtHCV IRESs into the Pol II non-continuous Tornado translation system, which contained a stop codon at the end of the split nLuc ORF (**Figure 4A**). We found that mutHCV IRES produced similar levels of luminescence as the wtHCV IRES (**Figure 4D**). Thus, the mutHCV IRES retained its ability to drive translation and can be used for continuous translation.

Next, we wanted to test the efficiency of the mutHCV continuous translation system. To test whether continuous translation was occurring, we cloned the mutHCV IRES into a continuous Tornado translation system where the IRES and split nLuc ORF contained no stop codons and a P2A sequence was included downstream of the IRES (**Figure 4A**). We then compared the protein expression from the mutHCV continuous translation system to the mutHCV non-continuous translation system. Interestingly, we saw only a 50% increase in luminescence by using a continuous translation system which suggests that the ribosome was only able to circumambulate ∼1-2 times on average (**Figure 4E**). This is likely due to the fact that the HCV IRES is highly structured, and structure is known to slow or even stop translation (Chen et al., 2013; Wen et al., 2008; Zheng et al., 2010).

We next considered the possibility that the slight increase in translational output from the mutHCV continuous translation system could express more protein than the CVB3 non-continuous translation system. To test this, we compared the translational output of the mutHCV continuous translation system to the CVB3 non-continuous translation system. The CVB3 non-continuous translation system produced ∼10-fold more luminescence than the continuous mutHCV translation system (**Figure 4E**). Thus, the mutHCV continuous translation system does not increase the protein output of the Tornado translation system.

Alternatively, we wanted to create a continuous translation system using an IRES that is less structured than a viral IRES. Recently, a high-throughput screen discovered thousands of endogenous IRES elements that drive translation of circular RNAs (Chen et al., 2021). These endogenous IRES elements drive translation using a small stem-loop structure, which is markedly less structure than the common viral IRESs (Chen et al., 2021). Moreover, many of the IRES elements identified in the screen are free of stop codons in at least one frame due to their 200 nt length. The combined benefit of having a simple structure and not requiring mutations may increase the ability of an endogenous IRES to drive continuous translation.

We first wanted to determine if an endogenous IRES could be used to drive continuous translation. To test this, we selected an IRES candidate from the screen (Chen et al., 2021) that exhibited high translational activity and lacked a stop codon in at least one frame—the *LIMA1* IRES. Second, we cloned the *LIMA1* IRES into a continuous or non-continuous Pol II-driven Tornado translation system using a split nLuc ORF and compared the translational output. Interestingly, we saw a ∼90-fold increase in luminescence from the *LIMA1* continuous translation system compared to the *LIMA1* non-continuous translation system (**Figure 4F**). This shows that endogenous IRESs may be the best candidates for continuous translation due to their short length and relatively unstructured nature.

Next, we wanted to compare the translational output from the *LIMA1* continuous translation system to the CVB3 non-continuous translation system. The *LIMA1* continuous translation system produced >15-fold less luminescence than the CVB3 non-continuous translation system (**Figure 4F**). Thus, even though continuous translation markedly enhances the protein output of the *LIMA1* IRES, its overall activity is still too low to express comparable amounts of protein to the CVB3 non-continuous system.

We next asked whether alternative endogenous IRES elements from the previously mentioned screen (Chen et al., 2021) would have higher translational output than the *LIMA1* IRES. To test this, we selected three additional endogenous IRESs (*CEP50, TGFBRG,* and *CHD2)* that exhibited high translational activity according to the screen and lacked a stop codon in at least one frame. We then cloned these IRESs into the continuous Tornado translation system using a split nLuc ORF and compared the translation output. We found that none of these endogenous IRESs produced more luminescence than the *LIMA1* continuous translation system (**Figure 4G**). Thus, endogenous IRES elements are unlikely to produce more protein than a viral IRES—despite being able to undergo continuous translation.

### The Tornado translation system provides lower, but more persistent levels of protein production than a linear mRNA translation system

We next wanted to compare the properties of circular mRNA, made with the CMV-CVB3 Tornado translation system, to a traditional linear cap-dependent mRNA.

First, we wanted to compare the overall protein expression level of the Tornado translation system to a linear cap-dependent mRNA translation system. To do this, we created a linear mRNA construct with the same split nLuc ORF as the Tornado translation system. The linear mRNA relies on cap-dependent translation (**Figure 5A**). We then compared the protein expression from the linear cap-dependent mRNA translation system to the Tornado translation system. We found that the Tornado translation system produced ∼5 fold less luminescence than the linear mRNA translation system (**Figure 5B**). Thus, a linear, cap-dependent translation system produces more protein than the Tornado translation system in a plasmid transfection setting.

We next asked if the lower amount of protein from the Tornado translation system compared to the linear mRNA translation system was due to the decreased protein output from CVB3-dependent translation compared to cap-dependent translation. To test this, we created an additional linear construct that relies on CVB3-dependent translation. The CVB3-dependent linear construct has the same split nLuc ORF and CVB3 IRES as the Tornado translation system, but in a linear form. To ensure the CVB3-dependent linear translation system would not undergo cap-dependent translation, we included an upstream ORF that ended with a stop codon (**Figure 5A**). We then compared the protein expression from the linear cap-dependent translation system to the linear CVB3-dependent translation system. We found that the linear CVB3-dependent translation system produced ∼10-fold less luminescence than the linear cap-dependent translation system (**Figure 5B**). Thus, CVB3-dependent translation leads to lower protein expression than cap-dependent translation. However, this could be due to decreased translational activity of the IRES or decreased RNA levels in transcripts that contain a CVB3 IRES.

We therefore wanted to determine whether the CVB3-dependent translation systems exhibited lower RNA expression than the cap-dependent translation system. To test this, we performed a northern blot using probes against the LgBiT on RNA from HEK293T cells transfected with the Tornado translation system, the linear CVB3-dependent translation system, and the linear cap-dependent translation system. We found that the linear cap-dependent translation system had ∼10-fold increased RNA expression compared to the linear CVB3-dependent transcript and ∼3-fold increased RNA expression compared to the Tornado translation system (**Figure 5C**). Moreover, when we normalized the amount of luminescence to the RNA expression, we found that the CVB3-dependent translation systems showed ∼3-fold lower luminescence (**Figure 5D**). Taken together, these data show that the decreased protein expression from the Tornado translation system compared to a linear cap-dependent translation system is due to the combination of decreased RNA expression of transcripts that contain the CVB3 IRES compared to cap-dependent transcripts and decreased translational activity of CVB3-dependent translation compared to cap-dependent translation.

Lastly, we wanted to determine if the Tornado translation system extended the duration of protein expression compared to a linear mRNA expression system. To test this, we used a tet-ON system to control transcription of the Tornado translation and linear mRNA expression systems. We generated stable cell lines that expressed the Tornado translation system, linear cap-dependent translation system, and linear CVB3-dependent translation system under a tetracycline-responsive promoter. We then pulsed the stable cell lines with tetracycline for 12 hours to induce transcription. Then, media was changed to tetracycline-free media. Cells were harvested at 0, 5, 10, 24, 48, and 72 hours later and assayed for RNA expression and luminescence. We found that the Tornado translation system provided ∼10-fold longer duration of luminescence and mRNA expression compared to both the linear cap-dependent and CVB3-dependent translation systems (**Figure 5E, 5F**). Thus, the Tornado translation system prolongs the duration of protein expression compared to a linear mRNA expression system encoding the same protein.

### The Tornado translation system can be used in multiple cell types

Next, we wanted to determine if the Tornado translation system can be used as a protein expression system in alternative cell types. We transfected HepG2 and ZR-75-1 cells with the Tornado translation system (CMV-CVB3) and detected luminescence in both these cell lines (**Figure S9A, B**). The Tornado translation system is therefore applicable to a variety of different cell types.

### The Tornado translation system can circularize long mRNAs

We next wanted to ensure that the Tornado system could circularize long mRNAs. The circular aptamers described in the original Tornado study were 100-350 nt long. In contrast, the split nLuc and CVB3 IRES together make a 1527 nt transcript. However, mRNAs can be much longer, so we wanted to determine if the Tornado system could be used to circularize the longer mRNAs. To test this, we generated a Tornado construct designed to express a circular mRNA that encodes for the spike protein from SARS-CoV-2 (Huang et al., 2020). This circular mRNA, including the CVB3 IRES, is 4719 nt long. We transfected HEK293T cells with a plasmid expressing the Tornado spike mRNA along with a plasmid expressing a linear spike RNA. We then quantified the RNA abundance after RNAse R treatment and found that the Tornado spike mRNA was markedly more resistant to RNAse R treatment than the linear spike mRNA (**Figure S10A**). The Tornado system can therefore be used to circularize long mRNAs.

### The Tornado translation system can be used to produce circular mRNA-containing VLPs

We next wanted to develop VLPs carrying circular mRNAs to achieve a longer duration of heterologous protein expression. To package circular mRNAs, we used the mRNA-containing lentiviral VLP system (Lu et al., 2019). This system comprises an envelope plasmid; a transfer plasmid that contains a gene of interest with an MS2 stem loop in its 3’UTR; and an integrase-deficient packaging plasmid that expresses MCP fused to the N-terminus of the nucleocapsid protein (Lu et al., 2019) (**Figure 6A**). We sought to modify this system to package circular mRNAs in VLPs. To do so, we created a transfer plasmid by cloning a MS2 stem loop into the 3’UTR of the CMV-CVB3 Tornado translation system that expresses an nLuc gene (**Figure 6A**). Since it contains the MS2 sequence, this RNA will be packaged into VLPs by binding to the MCP domain in the nucleocapsid protein.

**Figure 6.**
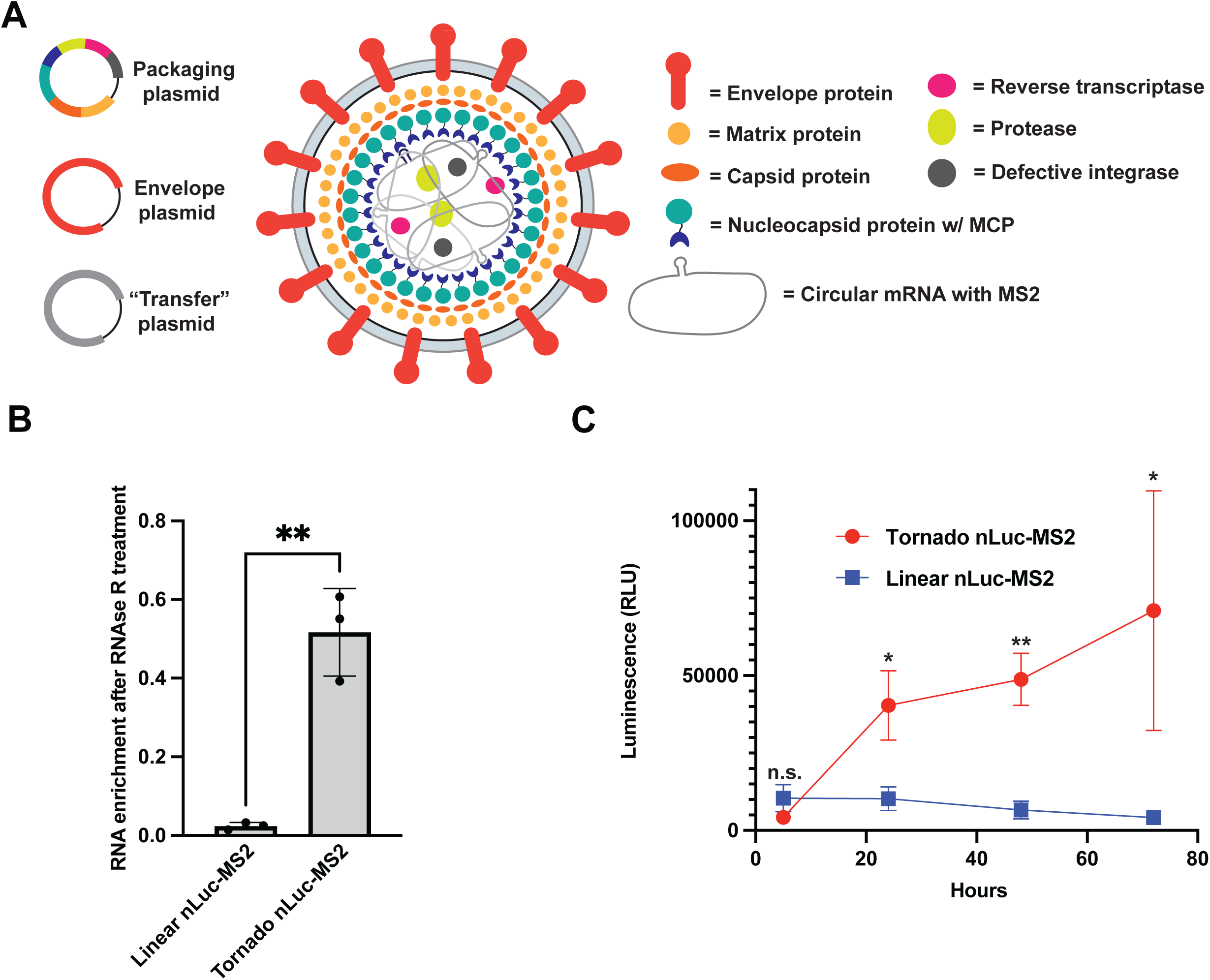
VLPs produced using the Tornado translation system exhibit increased level and duration of protein expression compared to conventional VLPs. (A) Schematic of the circular mRNA VLP system. The “transfer” plasmid comprises a Tornado translation system that expresses a circular mRNA with an MS2 stem loop. The packaging plasmid comprises the viral proteins required for VLP assembly including a nucleocapsid protein with a MCP domain fused to its N-terminus. The circular mRNA is packaged into the VLP by binding to the MCP domain that is fused to the N-terminus of the nucleocapsid protein. In this study, the envelope plasmid expresses a vesicular stomatitis virus G (VSV G) protein. However, the envelope plasmid can be exchanged for alternative envelope plasmids to achieve altered tropism. In this experiment, the mRNA being packaged with both the Tornado translation system and the linear cap-dependent mRNA expression system expresses an nLuc protein. (B) VLPs produced using the Tornado translation system contain circular mRNA. We extracted RNA from VLPs that were produced using the Tornado translation system (Tornado nLuc-MS2), or the linear mRNA expression system (Linear nLuc-MS2) as the transfer plasmid. RNA was treated with vehicle or RNase R to test whether the RNA is circular. RNA quantification was performed by qRT-PCR using primers that amplified a 126nt region of the nLuc gene (n = 3 biological replicates). Data are presented as mean values +/− one SD (n = 3 biological replicates). Significance was calculated using unpaired two-tailed student’s *t*-test. ****p<.0001, ***p<.001, **p<.01, *p<.05, n.s. p>.05. (C) VLPs that were produced using the Tornado translation system provide increased protein expression compared to VLPs produced by the linear mRNA expression system. We quantified luminescence from HEK293T cells transduced with VLPs that were produced using either the Tornado translation system (Tornado nLuc-MS2) (red) or the linear mRNA expression system (Linear nLuc-MS2) (blue) at hours 5, 24, 48, and 72 after transduction (n = 3 biological replicates). HEK293T cells were transduced at equal levels of VLP mRNA (**Figure S11A**). At 5 hours after transduction, cells transduced with VLPs that were produced using the linear mRNA expression system produced similar levels of luminescence to the cells transduced with VLPs that were produced using the Tornado translation system. At 24 hours after transduction and beyond, VLPs that were produced using the Tornado translation system produced >5-fold more luminescence expression than VLPs that were produced using the linear mRNA expression system. RLU = Relative Luminescence units. Data are presented as mean values +/− one SD (n = 3 biological replicates).

First, we wanted to ensure that we could package circular mRNAs into VLPs. Although the Tornado translation system expresses a predominantly circular mRNA (see **Figure 1C**), we wanted to rule out the possibility that linear precursors are packaged into the VLPs. To test this, we subjected RNA from VLPs produced using the Tornado translation system to RNase R treatment. As a control, we included a linear mRNA VLP system with the same nLuc ORF and MS2 sequence as our circular mRNA. We found that the viral RNA from VLPs packaged using the Tornado translation system were resistant to RNase R treatment while the viral RNA from the control VLPs made with linear RNAs were degraded (**Figure 6B**). Thus, the VLPs produced with the Tornado translation system primarily package circular mRNA.

### VLPs produced using the Tornado translation system exhibit increased level and duration of protein expression compared to conventional VLPs

We next wanted to determine whether VLPs produced using the Tornado translation system exhibit a longer duration of protein expression compared to VLPs produced using a linear mRNA expression system. To test this, we transduced HEK293T cells with equal amounts of infectious VLPs, as determined by the levels of nLuc mRNA in the VLP measured by qRT-PCR (**Figure S11A)**. At the initial 5-hour time point, the cells transduced with the VLPs produced using the Tornado translation system produced a similar amount of luminescence as the cells transduced with VLPs produced using a linear mRNA expression system (**Figure 6C**). However, at 24 hours after transduction and beyond, VLPs that were produced using the Tornado translation system produced >5-fold more luminescence than VLPs that were produced using the linear mRNA expression system (**Figure 6C**). This result is consistent with the longer half-life of circular mRNA, which would allow prolonged synthesis and accumulation of protein over time. We also asked if these VLPs can be used in other cell types. We therefore tested SH-SY5Y neuroblastoma cells, which similarly showed a ∼5-fold increase in luminescence by using a VLP produced using the Tornado translation system compared to a VLP produces using linear mRNA expression system (**Figure S12A**). Overall, these data show that VLPs produced using the Tornado translation system enable extended duration of protein expression and higher overall protein expression compared to VLPs produced using a linear mRNA expression system.

## DISCUSSION

Circular mRNA is an important type of mRNA therapeutic due to its extended half-life in the mammalian cytosol. However current VLP approaches are only capable of packaging and delivering linear mRNAs. Here we describe the development of VLPs with circular mRNAs, thus enabling VLPs to exert longer duration of protein expression, and in some cases, increased overall protein expression, compared to VLPs with linear mRNAs. This approach is enabled by the Tornado RNA circularization approach, in which RNAs are transcribed with a transcript of interest flanked by specific ribozymes. After autocatalytic RNA cleavage, the resulting RNA ends are compatible for ligation with RtcB. Through several different types of optimizations, we identify altered sequence requirements for the Tornado RNA, as well as specific promoters and IRESs that result in a translatable circular mRNAs that are efficiently packaged into VLPs. Using this approach, we show that the VLPs produced using the Tornado translation system can increase the duration and level of protein expression, thus demonstrating the potential use of this technology for mRNA delivery.

As part of our study, we examined circular mRNA generated by the backsplicing system. This method generates circular RNA by using the molecular mechanism that occurs with the endogenous genes, such as *ZKSCAN1*. In this pathway, intronic sequences have sequence complementarity, which thus leads to intron-intron interactions that subsequently facilitate rare backsplicing events (Liang and Wilusz, 2014). However, our experiments showed that the predominant product of this reaction was a linear product, likely reflecting the forward splicing reaction. Other groups similarly found that the backsplicing system generated linear forward splicing products (Ho-Xuan et al., 2020; Jiang et al., 2021). In contrast, the Tornado system readily generated large circular mRNAs (up to 4719 bp), with minimal detectable linear precursors. This reflects the highly efficient nature of circularization using the Tornado approach. Thus, for experiments requiring generation of small or large RNA circles in cells, the Tornado approach should be used.

The VLP system has the benefit that it can be pseudotyped to enable cell-type specific infection (Cronin et al., 2005; Hamilton et al., 2021; Naldini et al., 1996). Notably, recent variants of the VLP system have been described using endogenous retrotransposons, termed the “SEND” (selective endogenous encapsidation for cellular delivery) system (Segel et al., 2021). This approach may be particularly useful for delivering mRNA without unwanted immune effects that would presumably occur using repeated dosing of current VLPs.

Our current system relies on circular mRNAs made using the CMV promoter, which is a Pol II promoter. However, we found that much more circular mRNA was generated using a U6 promoter, which is a Pol III promoter. Thus, even though Pol III normally generates small RNAs such as tRNAs, it can be used to make large circular mRNAs. However, we needed to mutate the IRES to remove Pol III termination elements, which in turn reduced the activity of the IRES. It will be important to develop Pol III-compatible IRESs that are also highly efficient for translation initiation to take advantage of the high RNA expression seen with the Pol III system.

We also compared different IRESs, including natural endogenous IRESs that were recently found in endogenous circular mRNAs (Chen et al., 2021). Our assay for measuring IRES activity does not have the drawback that is commonly seen in the classic bicistronic IRES reporter assay (Harger and Dinman, 2003) in which an IRES and reporter gene is placed in the 3’UTR of a gene. In this assay, any translation of the reporter is presumed to reflect IRES-mediated translation rather than cap-dependent translation. However, this assay has been criticized because cryptic transcription initiation sites can produce transcripts that initiate in the 3’UTR region, thus providing cap-dependent translation of the reporter (Kozak, 2003, 2005). As a result, a sequence may be falsely thought to function as an IRES when it instead functions as an alternative transcription initiation site. Our assay solves this problem. In our assay, the reporter is only functional if the RNA has circularized, which only occurs on transcripts that are transcribed from the intended promoter. Any cryptic transcription initiation would not lead to a circle since the 5’ ribozyme would not be present. In this way, we can very accurately measure the activity of any IRES. Our data show that the translational activity of the HRV-B3 IRES, recently tested in a large-scale screen of novel IRESs (Chen et al., 2022), is similar to the CVB3 IRES. We also found that the endogenous IRESs recently discovered in a high-throughput screen (Chen et al., 2021), are several orders of magnitude less effective than the CVB3 IRES, although we were able to detect IRES activity from at least one of the tested endogenous IRESs. Overall, the approach described here can be useful for screening and testing new IRESs.

Although our study emphasizes the use of circular mRNA for improving VLPs, it should be noted that cellular expression of circular mRNAs can be valuable for other applications. For example, plasmid therapeutics rely on the expression of the encoded mRNAs, but the plasmid DNA is often epigenetically silenced (Chen et al., 2004), limiting the duration of action of the plasmid DNA therapeutic. Adenoviral vectors similarly deliver a DNA that is readily silenced, thus limiting the duration of protein expression from adenoviral vectors (Brooks et al., 2004). Plasmid and adenoviral vector-based therapeutics could have a longer duration of action if they expressed circular mRNAs using the Tornado translation system.

### SIGNIFICANCE

We developed a method for expressing circular mRNAs in mammalian cells—the Tornado translation system. The Tornado translation system markedly increases the amount of circular mRNA that can be made in cells compared to the previous system, the backsplicing system. In a series of experiments, we selected the internal ribosomal entry site (IRES) and promoter that lead to high-level protein output from the Tornado translation system. We then showed that the Tornado translation system can be used to produce virus-like particles (VLPs) that exhibit increased protein expression compared to VLPs that are produced using a standard linear mRNA expression system. Overall, we show that genetically encoding circular mRNAs can improve the therapeutic potential of VLPs and potentially other modalities such as plasmid and adenoviral vector-based therapeutics.

### METHOD DETAILS

#### Reagents

Unless otherwise stated, all reagents were from Sigma-Aldrich except for cell culture reagents, which were from Invitrogen. These reagents were used without further purification.

#### Tornado stem design

The Tornado circularization junction stem was designed using mfold (http://www.unafold.org/mfold/applications/rna-folding-form.php). Stems were designed to have bulges every ∼10 base pairs.

#### Split nLuc design

Split nLuc ORF was codon optimized to avoid Pol III termination signals using the codon optimization tool from Integrated DNA Technologies. The frame of the Tornado circularization stem was chosen to avoid stop codons. To improve the sensitivity of the protein readout, the split nLuc contained a C-terminal 2x Glutamine degron.

#### Cloning Pol III transcripts

Split nLuc, IRES, and partial Tornado sequences were chemically synthesized as gene blocks (Integrated DNA Technologies), then cloned into the NotI and SacII sites of the pAV-U6+27-Tornado-Broccoli plasmid (Addgene #124360). Split nLuc ORF was codon optimized to be compatible with a Pol III promoter. Changing the IRES was done by cloning a gene block into the EcoRI and BsiWI internal restriction sites. All plasmids were sequenced (Psomagen) to verify identity.

#### Cloning Pol II transcripts

Split nLuc, nLuc, spike, IRES, and Tornado sequences were chemically synthesized as gene blocks (Integrated DNA Technologies). Gene blocks were cloned into the BamHI and XhoI sites of pcDNA3.1+ vector backbone. Changing the IRES was done by cloning a gene block into the EcoRI and BsiWI internal restriction sites. Minor alterations such as stop codon insertion or IRES point mutations were done using a QuikChange Site-directed mutagenesis kit II (Agilent) according to manufacturer’s instructions. All plasmids were sequenced (Psomagen) to verify identity.

#### Cloning of backsplicing system sequences

A gene block containing the same exact sequence as the CMV-CVB3 Tornado translation system was synthesized (Integrated DNA Technologies) and cloned into the EcoRV and SacII restriction site of the pcDNA3.1(+) CircRNA Mini Vector (Addgene #60648). The Tornado circularization stems were included at the 5’ end of the sequence to ensure the ORF from the Tornado Translation system would match the ORF from the backsplicing-based system. After cloning, this plasmid was sequenced (Psomagen) to verify identity.

#### Cell culture and transfection

HepG2 (ATCC HB-8065) and HEK293T (ATCC CRL-11268) cells were cultured with ×1 DMEM (ThermoFisher #11995-065) with 10% fetal bovine serum (FBS), 100 U/ml penicillin and 100 μg/ml streptomycin under standard tissue culture conditions. ZR-75 (ATCC CRL-1500) were cultured using RPMI 1640 Medium with no phenol red (ThermoFisher #11835030) with 10% fetal bovine serum (FBS), 100 U/ml penicillin and 100 μg/ml streptomycin under standard tissue culture conditions. SH-SY5Y were cultured with F-12/DMEM (ThermoFisher #11320033), 20% fetal bovine serum (FBS), 100 U/ml penicillin and 100 μg/ml streptomycin. Cells were cultured at 37 °C and 5% CO2 and passaged every 2–3 days. TrypLE Express (ThermoFisher #12604013) was used to lift cells for passaging. Cells were plated at a density of 2x10^4^ cells/cm^2^ 20 hours before transfection. Cells were transfected using a 3:1 ratio of FuGENE (Promega #E5911) to DNA in OptiMEM I Reduced Serum Media (ThermoFisher, #31985). Unless otherwise stated, all cells were harvested 72 hours after transfection for downstream applications.

#### Protein expression analysis

Cells were harvested by directly lifting cells with ×1 phosphate buffered saline (PBS) (ThermoFisher, no. 10010031). For luminescence assays, cells were harvested 72 hours after transfection unless otherwise stated. Media was aspirated off cells then cells were resuspended in PBS. 50µl of cell suspension was transferred to a flat-bottomed white-walled 96-well plate (Corning). Nano-Glo Luciferase Assay System (Promega #N1110) reagent was prepared according to manufacturer’s instructions. 50ul of Nano-Glo reagent was added to each well of cell suspensions. The plate was gently shaken, then luminescence detection was done using SpectraMax iD3 (Molecular Devices) machine with SoftMax Pro (v.7.1) software using the luminescence acquisition settings (Endpoint luminescence, 96 Well Standard opaque plate, integration time 1000 ms, 1 mm read height).

#### RNA extraction

RNA was harvested from cultured cells by removing media and detaching enzymatically or directly lifting cells with ×1 phosphate buffered saline (PBS) (ThermoFisher, #10010031). Cell suspensions were mixed with TRIzol LS Reagent (Invitrogen, no. 10296010), then frozen and stored at –20 °C or purified immediately according to the manufacturer’s instructions.

#### RNase R reactions

Following RNA extraction, RNA concentration was quantified using a NanoDrop 2000 (Thermo Scientific). Equal concentrations of RNA were added to two tubes and were treated with RNase R (Biosearch Technologies RNR07250) according to manufacturer’s instructions. Following RNase R reaction, RNA was purified using the RNA Clean & Concentrator kit (Zymo Research, no. R1015).

#### Northern blot

Following RNA extraction or RNase R reactions, RNA was blotted using the NorthernMax kit (ThermoFisher #AM1940) according to manufacturer’s instructions. Antisense DNA probes designed to bind to either the LgBiT or *ZKSCAN1* exon2/3 were synthesized with a 5’Biotin (Integrated DNA Technologies). Band detection was done using the Chemiluminescent Nucleic Acid Detection Module (ThermoFisher #89880) according to manufacturer’s instructions. The blot was imaged using ChemiDoc MP imager (Bio-Rad) with the chemiluminescent band detection setting. RNA quantification from northern blots was done using Image Lab (v.5.2.1) software.

#### qRT-PCR for split nLuc

Following RNA extraction or RNase R reactions, RNA was treated with DNase (ThermoFisher #EN0521) according to the manufacturer’s instructions. RNA was then directly used for cDNA synthesis with the Superscript III kit (ThermoFisher #12574026). cDNA was diluted 1:10 added to Eppendorf twin.tec 96 *real-time* PCR Plate (Eppendorf #0030132700) along with iQ Syber Green Supermix (Bio-Rad #1708880) and primers. Primers designed to amplify a 150nt region in the LgBiT region and reference primers for *GAPDH* were chemically synthesized (Integrated DNA Technologies). qPCR was done using the Eppendorf Realplex qPCR machine. RNA quantification was done by using the 2^-[ΔCt(target)-ΔCt(reference)]^ method.

#### NCBI BLAST searches

EMCV IRES sequence was aligned against viral (taxid:10239) reference genomes (refseq_genomes) with the somewhat similar (blastn) program on NCBI BLAST (https://blast.ncbi.nlm.nih.gov/Blast.cgi).

#### RNA/protein half-life experiments

Flip-In-293 cells (ThermoFisher #R75007) were cultured with ×1 DMEM (ThermoFisher #11995-065) with 10% fetal bovine serum (FBS), 100 U/ml penicillin and 100 μg/ml streptomycin under standard tissue culture conditions. Cell lines that stably express the Tornado (CMV-CVB3), Linear (CVB3), and Linear (Cap) mRNAs were made by using the Flp-In T-Rex Core Kit (ThermoFisher #K6500-01). Cells were plated at a density of 2x10^4^ cells/cm^2^ 16 hours before tetracycline treatment. Cells were then treated with 1 μg/mL tetracycline (Santa Cruz Biotechnology) for 12 hours then the media was changed to tetracycline-free media. Cells were harvested at 0, 5, 10, 24, 48, and 72 hours after tetracycline withdrawal. Cells were harvested by directly lifting cells with ×1 phosphate buffered saline (PBS) (ThermoFisher #10010031) then split in half for both protein quantification and RNA quantification.

qRT-PCR RNA quantification normalizes the target RNA level to the reference RNA level. This means the level of tet-inducible RNA expression will appear to go down over time as the HEK293 cells divide and make more *GAPDH* while the tet-inducible gene is no longer being expressed. RNA quantification was therefore adjusted to the cell count at every time point. Cell counts were obtained using Countess 3 automated cell counter (ThermoFisher). In addition, the RNA expression level needs to be normalized to the level of RNA expressed when no tetracycline is added. The equation for this normalization is as follows:

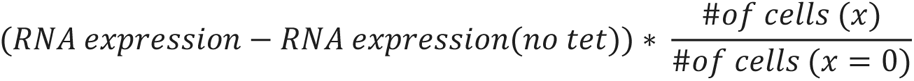

Where x is the hour that the RNA was harvested, and *RNA expression* was calculated using the **qRT-PCR for split nLuc** method.

#### VLP production

MCP-modified packaging plasmid pspAX2-D64-NC-MCP (Addgene #122944), VSVg envelope plasmid pMD2.G (Addgene #12259), and transfer plasmid Tornado/Linear nLuc-MS2 were transfected into 80% confluent HEK293T cells at a ratio of 3:1.5:4.5. 3 μg of total DNA was transfected into each well of a 6-well plate. Cells were transfected using a 3:1 ratio of FuGENE (Promega #E5911) to DNA in OptiMEM I Reduced Serum Media (ThermoFisher, #31985).

Media was changed 24 hours after transfection. VLPs were harvested and filtered through a 45-micron filter three days after transfection.

#### VLP transduction

Unconcentrated VLPs were diluted according to their viral RNA titers (see “**RNA isolation from VLPs and RT-qPCR analysis**”) in fresh media and added to 5 x 10^4^ cells in a 12-well plate. At the first collection time, cells were washed with 1ml of ×1 phosphate buffered saline (PBS) (ThermoFisher #10010031) before harvesting. Media for subsequent time points was replaced with fresh media after the first collection time point. At each collection time point, cells were harvested and subject to protein expression analysis (see “**Protein expression analysis**” section).

#### RNA isolation from VLPs and RT-qPCR analysis

RNA from VLPs was extracted using the QIAmp Viral RNA Mini Kit (Qiagen #52904) according to manufacturer’s instructions. RNA was then treated with RNase R (See RNase R reactions section) then DNAse (ThermoFisher #EN0521) according to the manufacturer’s instructions. RNA was then directly used for cDNA synthesis with the Superscript III kit (ThermoFisher #12574026). cDNA was diluted 1:10 added to Eppendorf twin.tec 96 *real-time* PCR Plate (Eppendorf #0030132700) along with iQ Syber Green Supermix (Bio-Rad #1708880) and primers. Primers that amplified a 125nt amplicon within the nLuc gene were designed using Integrated DNA technologies. RNA abundance was calculated by using the 2^-[ΔCt(target)-ΔCt(reference)]^ equation with the reference being a no RNA control. For RNase R reactions, RNA was quantified using the 2^-(Ct^ ^+RNase^ ^R−Ct-RNase^ ^R)^ equation.

## QUANTIFICATION AND STATISTICAL ANALYSIS

All data are expressed as means ± SD with the number of independent experiments (n) listed for each experiment. Statistical analyses were performed using Excel (Microsoft) and Prism (GraphPad).

### DATA AND CODE AVAILABILITY

Tornado translation expression plasmids will be made available on Addgene. All gene blocks used in the paper will be available in the supplement.

## ACKNOWLEDGMENTS

We thank J. L. Litke for his comments and suggestions. This work was supported by NIH grant R35NS111631 (to S.R.J.) and NIH fellowship no. F31NS125945 (to M.J.U.).

## AUTHOR CONTRIBUTIONS

S.R.J. and M.J.U. conceived and designed the experiments. M.J.U. performed the experiments and analyzed the data. S.R.J. and M.J.U. wrote the manuscript.

## Supplemental Information

**Figure S1, related to Figure 1.**
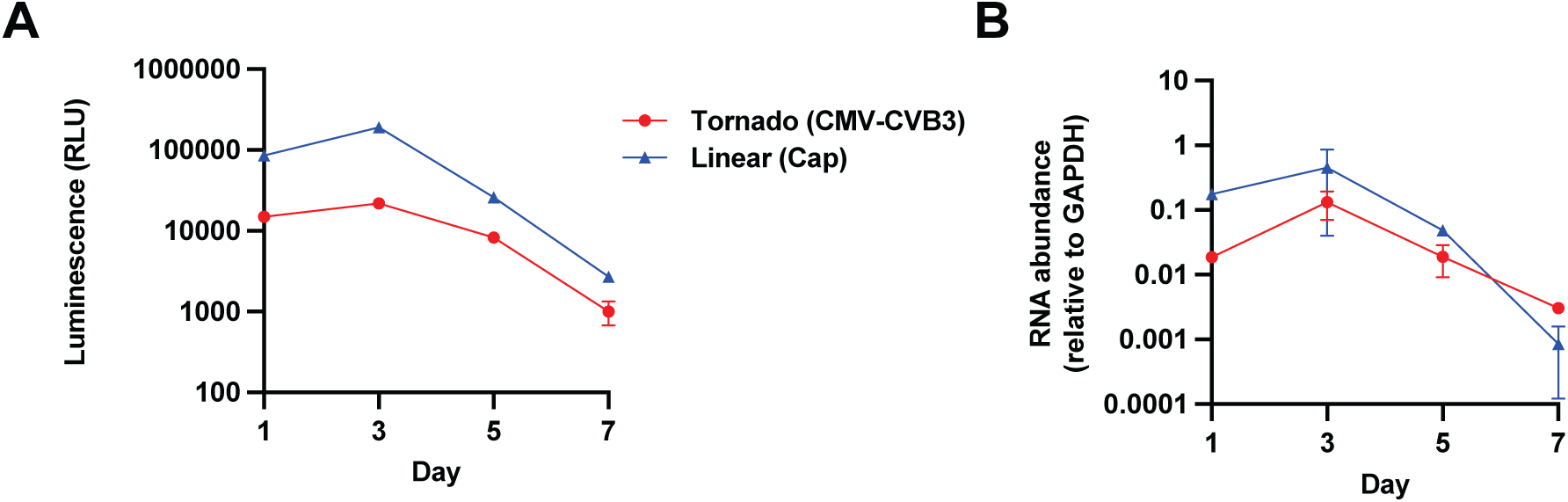
Peak protein and mRNA expression is reached at day 3 after transfection for the Tornado translation system and the linear cap-dependent mRNA expression system. (A) Peak protein expression is reached at day 3 for the Tornado translation system and the linear cap-dependent mRNA expression system. We quantified luminescence at days 1, 3, 5, and 7 after HEK293T cells were transfected with plasmids expressing the Tornado translation system (Tornado (CMV-CVB3)) and the linear cap-dependent mRNA expression system (Linear (Cap)). Peak luminescence from the Tornado translation system and the linear cap-dependent mRNA expression system is reached at day 3. RLU = Relative Luminescence units. Data are presented as mean values +/− one SD (n = 3 technical replicates). (B) Peak RNA expression is reached at day 3 for the Tornado translation system and the linear cap-dependent mRNA expression system. We quantified RNA at days 1, 3, 5, and 7 after HEK293T cells were transfected with plasmids expressing Tornado translation system (Tornado (CMV-CVB3)) and the linear cap-dependent mRNA expression system (Linear (Cap)) by performing qRT-PCR with primers that amplify a 120 nt region of the LgBit. We used *GAPDH* to normalize RNA abundance. Peak RNA abundance from the Tornado translation system and the linear cap-dependent mRNA expression system is reached at day 3. Data are presented as mean values +/− one SD (n = 2 technical replicates).

**Figure S2, related to Figure 1.**
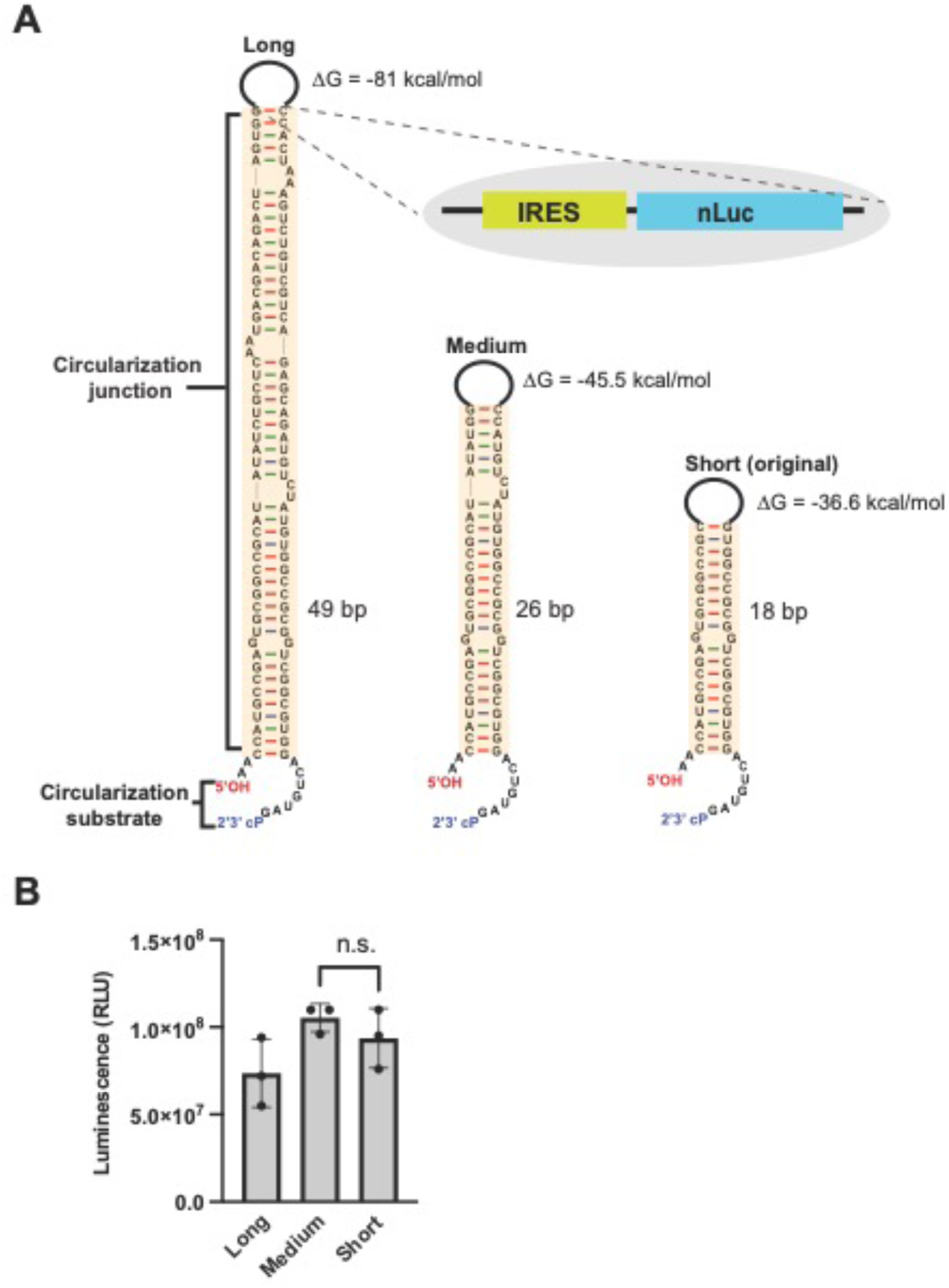
Lengthening the circularization junction stem does not increase protein output from the Tornado translation system. (A) mFold representation of three Tornado circularization junction stems with various lengths. The stems are designed to flank the IRES (CVB3) and nLuc sequence to facilitate circularization. More efficient circularization is reflected by increased luminescence. (B) Lengthening the circularization junction stem does not increase protein output from the Tornado translation system. We quantified luminescence from HEK293T cells transfected with plasmids carrying the Tornado translation system with the short, medium, and large stems (n = 3 biological replicates). All three constructs were expressed using the CMV promoter and used the CVB3 IRES to drive translation of the nLuc mRNA. The three stems produced similar levels of luminescence. RLU = Relative Luminescence units. Data are presented as mean values +/− one SD (n = 3 biological replicates). Significance was calculated using unpaired two-tailed student’s *t*-test. ****p<.0001, ***p<.001, **p<.01, *p<.05, n.s. p>.05.

**Figure S3, related to Figure 2.**
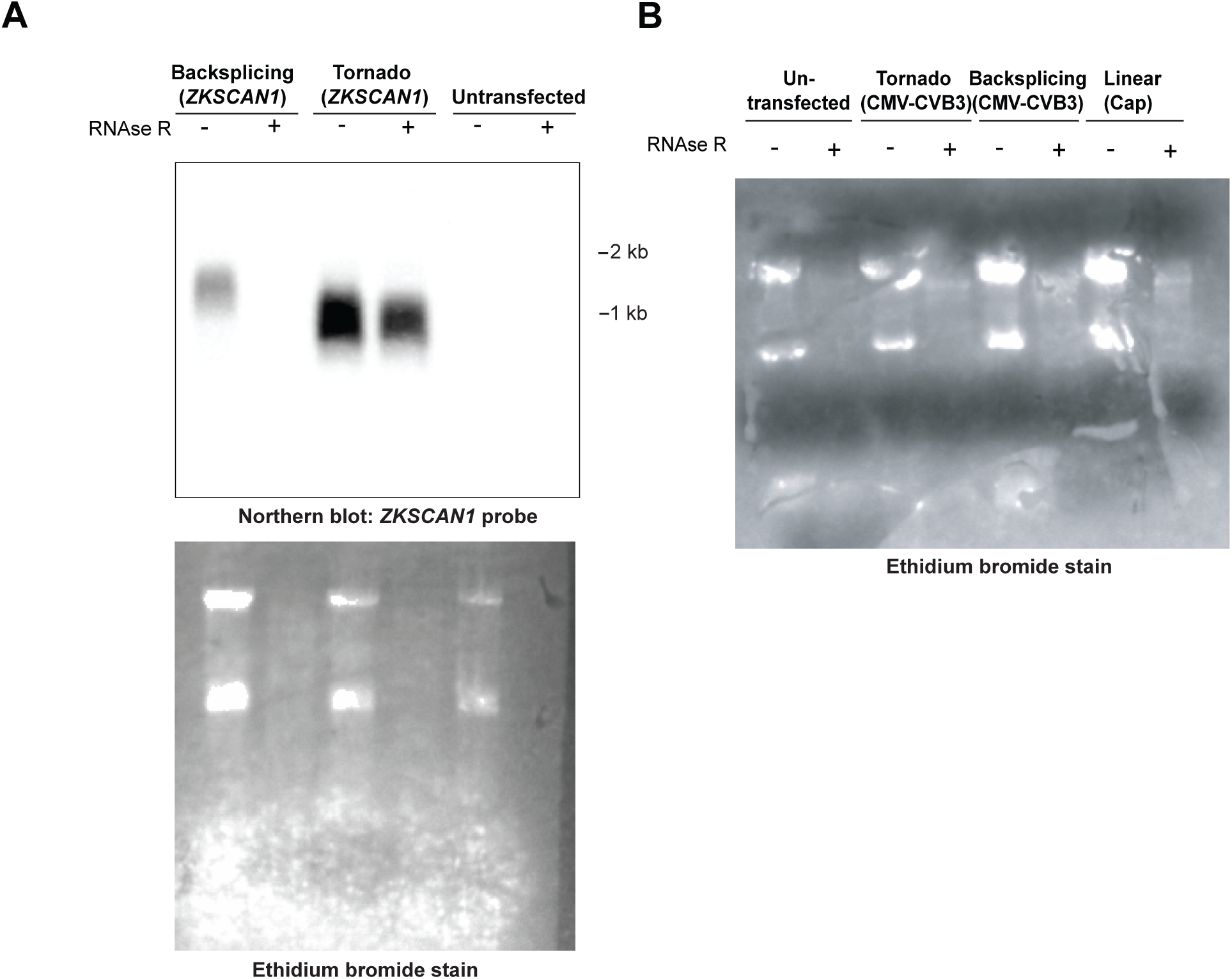
The Tornado translation system produces more circular RNA than the backsplicing system. (A) The Tornado translation system produces more circular RNA than the backsplicing system. HEK293T cells were transfected with plasmids expressing the Tornado translation system and backsplicing system containing a *ZKSCAN1* exon 2/3 insert. RNA was treated with vehicle or RNase R to test whether the RNA is circular. The Tornado translation system produces an RNA that is primarily in circular form. The backsplicing system produces an RNA that is primarily in a linear form. Ethidium bromide-stained blot shows successful RNase R treatment. (B) An ethidium bromide stain of the northern blot shown in Figure 2C. Disappearance of rRNA bands in RNase R treated samples shows successful RNase R treatment.

**Figure S4, related to Figure 3.**
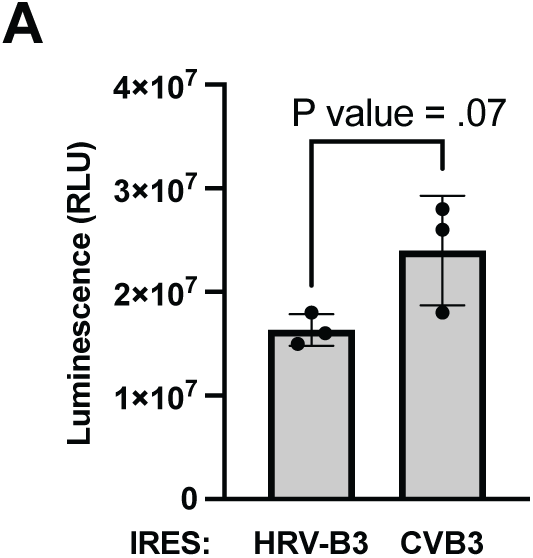
The CVB3 IRES produces a similar amount of protein as the HRV-B3 IRES. (A) The CVB3 IRES produces a similar amount of protein as the HRV-B3 IRES. We quantified luminescence from HEK293T cells transfected with plasmids expressing the CVB3 or HRV-B3 IRES. Both constructs were expressed using a Pol II-driven (CMV) Tornado translation system with the split nLuc ORF. The CVB3 and HRV-B3 IRES produce similar levels of luminescence. Using a P value cutoff of <.1, the CVB3 IRES produced more luminescence than the HRV-B3 IRES. RLU = Relative Luminescence units. Data are presented as mean values +/− one SD (n = 3 biological replicates). Significance was calculated using unpaired two-tailed student’s *t*-test.

**Figure S5, related to Figure 3.**
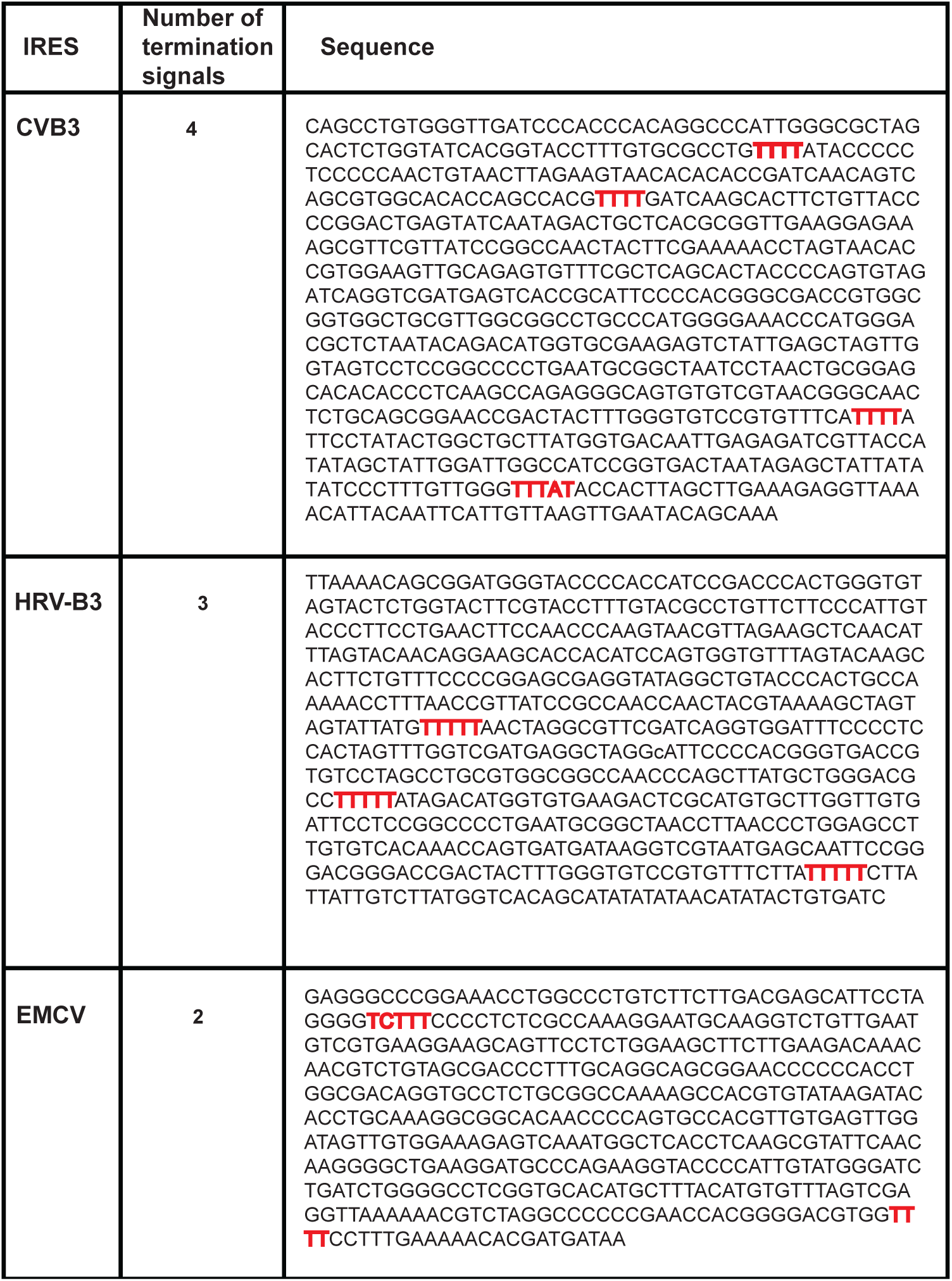
Visualization of Pol III termination elements in CVB3, EMCV and HRV-B3. (A) Sequences of CVB3, EMCV, and HRV-B3 and their Pol III termination elements. CVB3 has four Pol III termination elements. HRV-B3 has three Pol III termination elements. EMCV has two Pol III termination elements.

**Figure S6, related to Figure 3.**
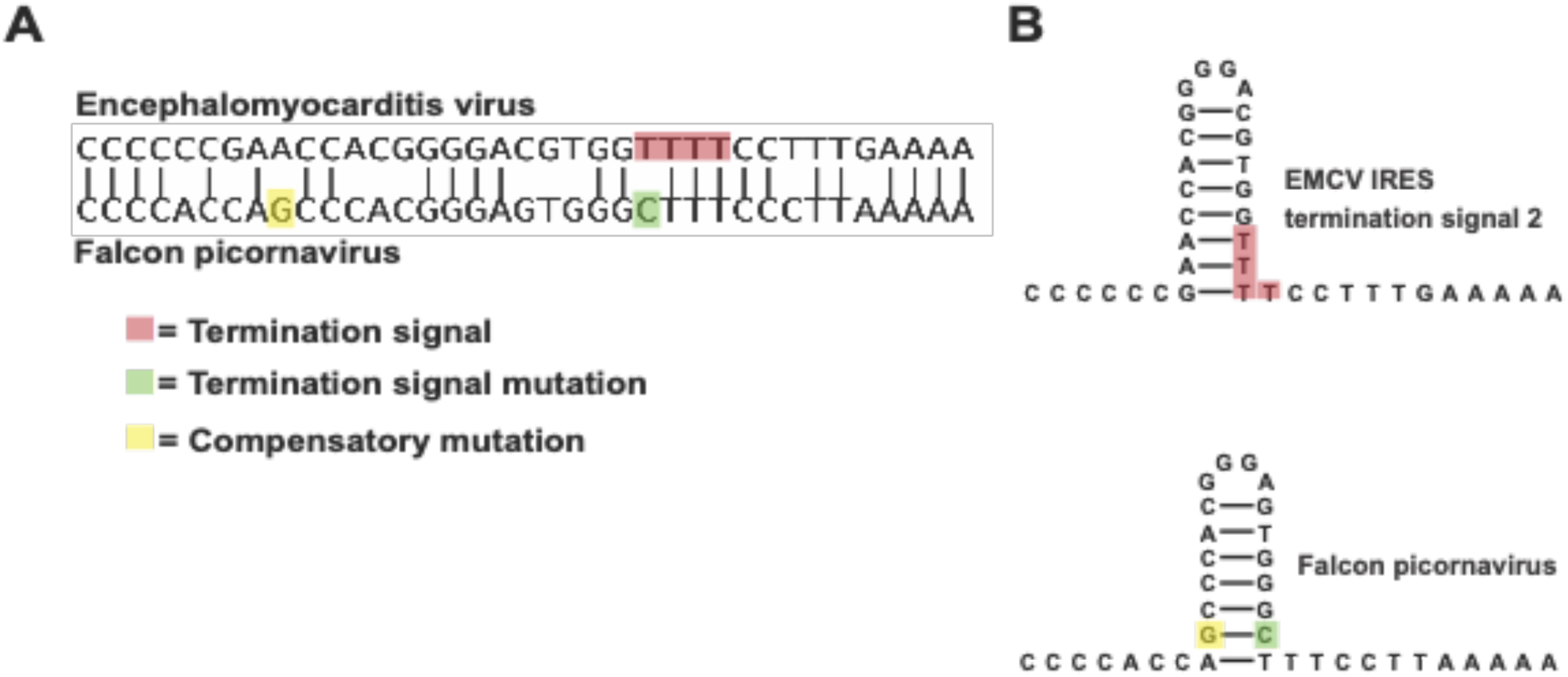
Falcon picornavirus IRES can guide the EMCV termination signal 2 mutation. (A) The falcon picornavirus IRES shares a similar sequence as the EMCV IRES but does not contain a Pol III termination signal. BLAST alignment shows sequence similarity between the EMCV IRES and the falcon picornavirus IRES. The red box shows the EMCV termination signal 2. The green box shows the mutation that eliminates the Pol III termination signal in the falcon picornavirus. The yellow box shows the compensatory mutation that maintains the structure of the IRES without the Pol III termination element. (B) mFold structural predictions show that the falcon picornavirus maintains the structure of the stem loop that contains the EMCV termination signal 2 mutation but does not have a Pol III termination signal.

**Figure S7, related to Figure 3.**
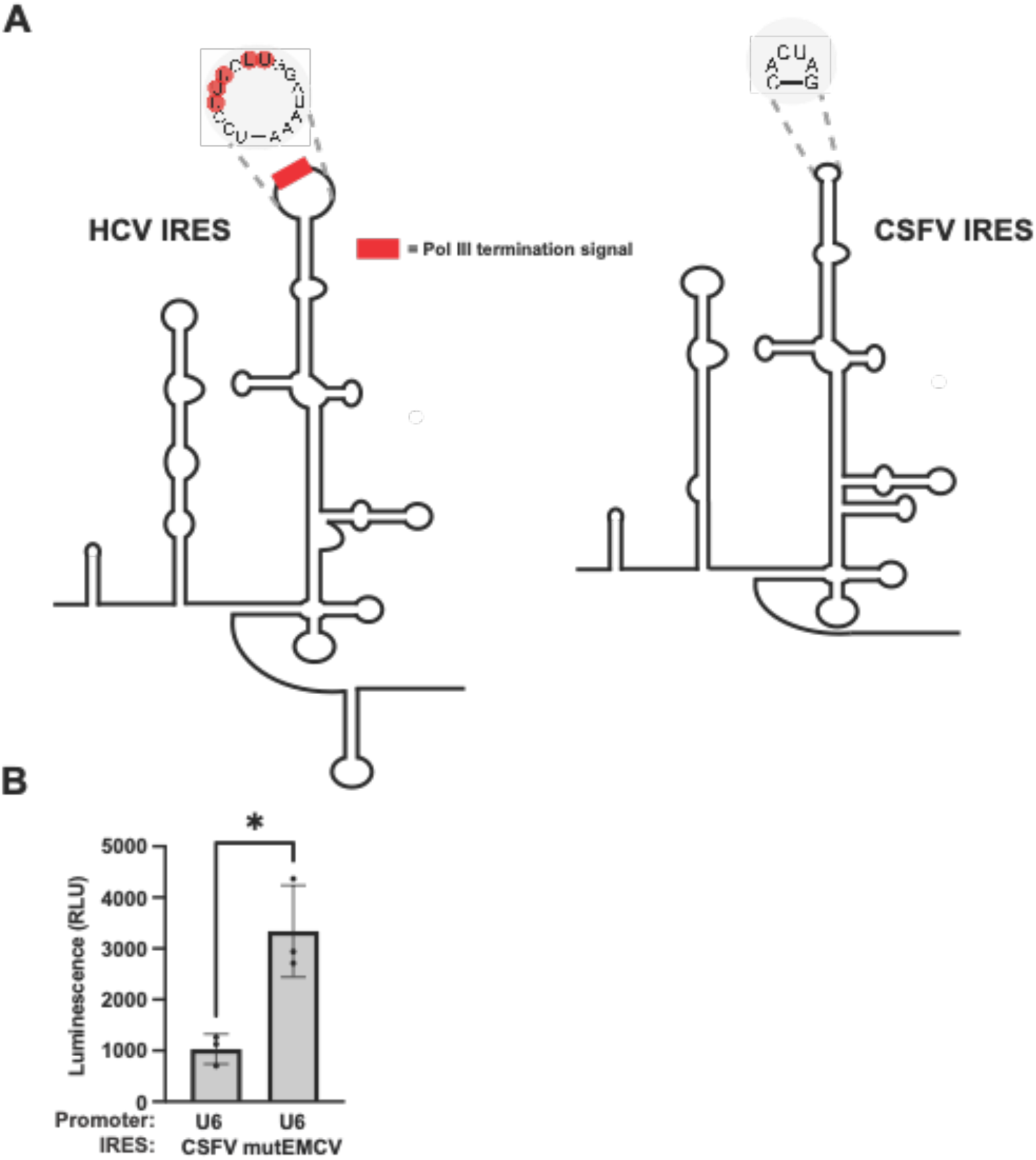
The Pol III-compatible mutHCV IRES produces more protein than the naturally Pol III-compatible CSFV IRES. (A) HCV IRES and CSFV IRES are similar in structure, but not sequence. CSFV has a similar structure as the HCV IRES yet lacks the Pol III termination element that is present in HCV. (B) mutHCV produces more protein than the CSFV in a Pol III-driven Tornado translation system. We quantified luminescence from HEK293T cells transfected with plasmids expressing the CSFV (U6 CSFV) and mutEMCV (U6 mutEMCV) IRES. Both constructs were expressed using a Pol III-driven (U6) Tornado translation system with the split nLuc ORF. The CSFV IRES produced 3-fold less luminescence than the mutEMCV IRES. RLU = Relative Luminescence units. Data are presented as mean values +/− one SD (n = 3 biological replicates). Significance was calculated using unpaired two-tailed student’s *t*-test. ****p<.0001, ***p<.001, **p<.01, *p<.05, n.s. p>.05

**Figure S8, related to Figure 1, Figure 3, and Figure 5.**
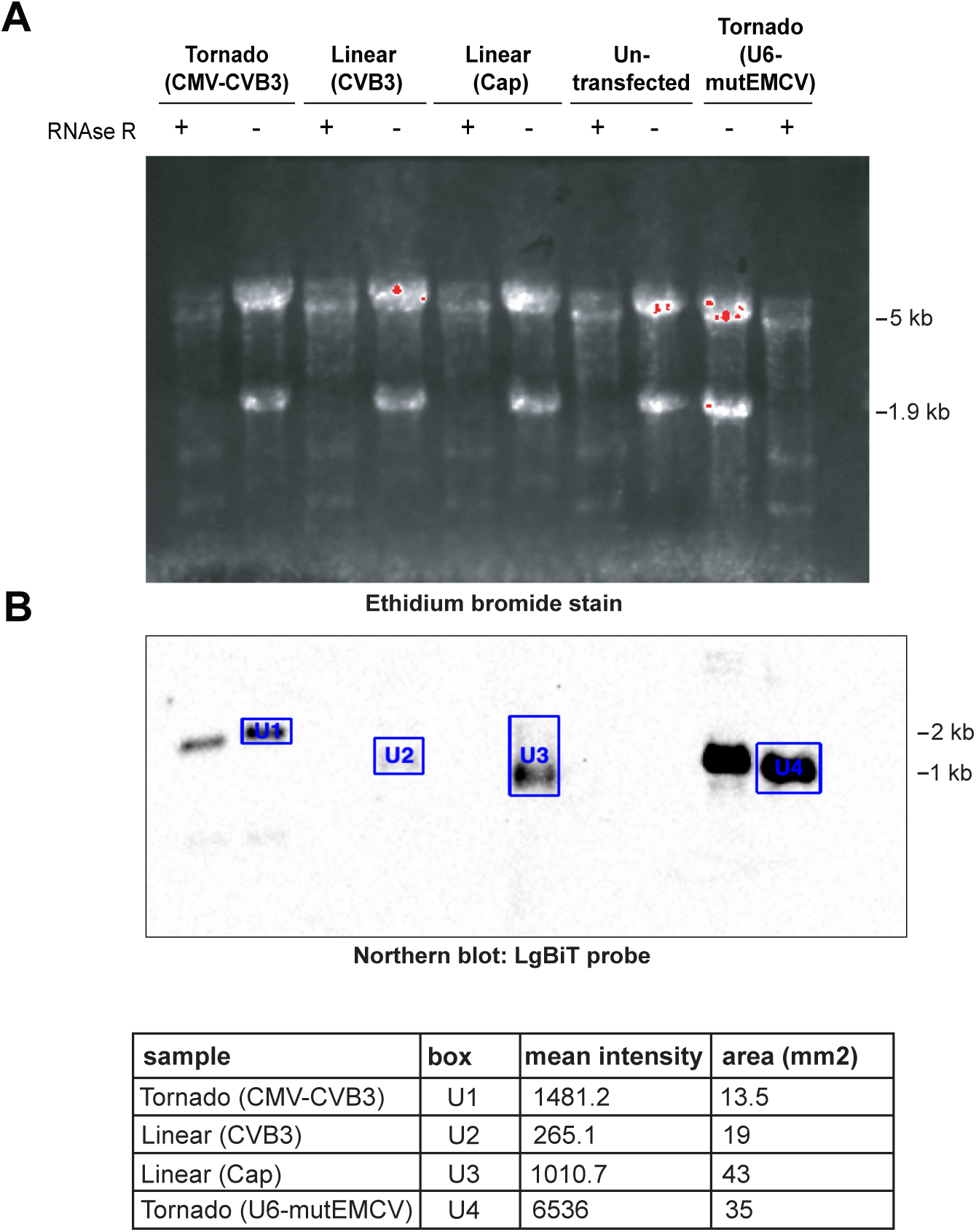
Northern blot shows successful RNase R treatment and can be used to quantify RNA expression levels. (A) Ethidium bromide stain of northern blot shown in **Figure 1C**, **3E, and 5C**. Disappearance of rRNA bands in RNase R treated samples shows successful RNAse R treatment. Ethidium bromide stain shows similar loading of RNA samples. (B) RNA quantification from the northern blot. Quantification of RNA was done by multiplying the mean intensity by the area of the band.

**Figure S9.**
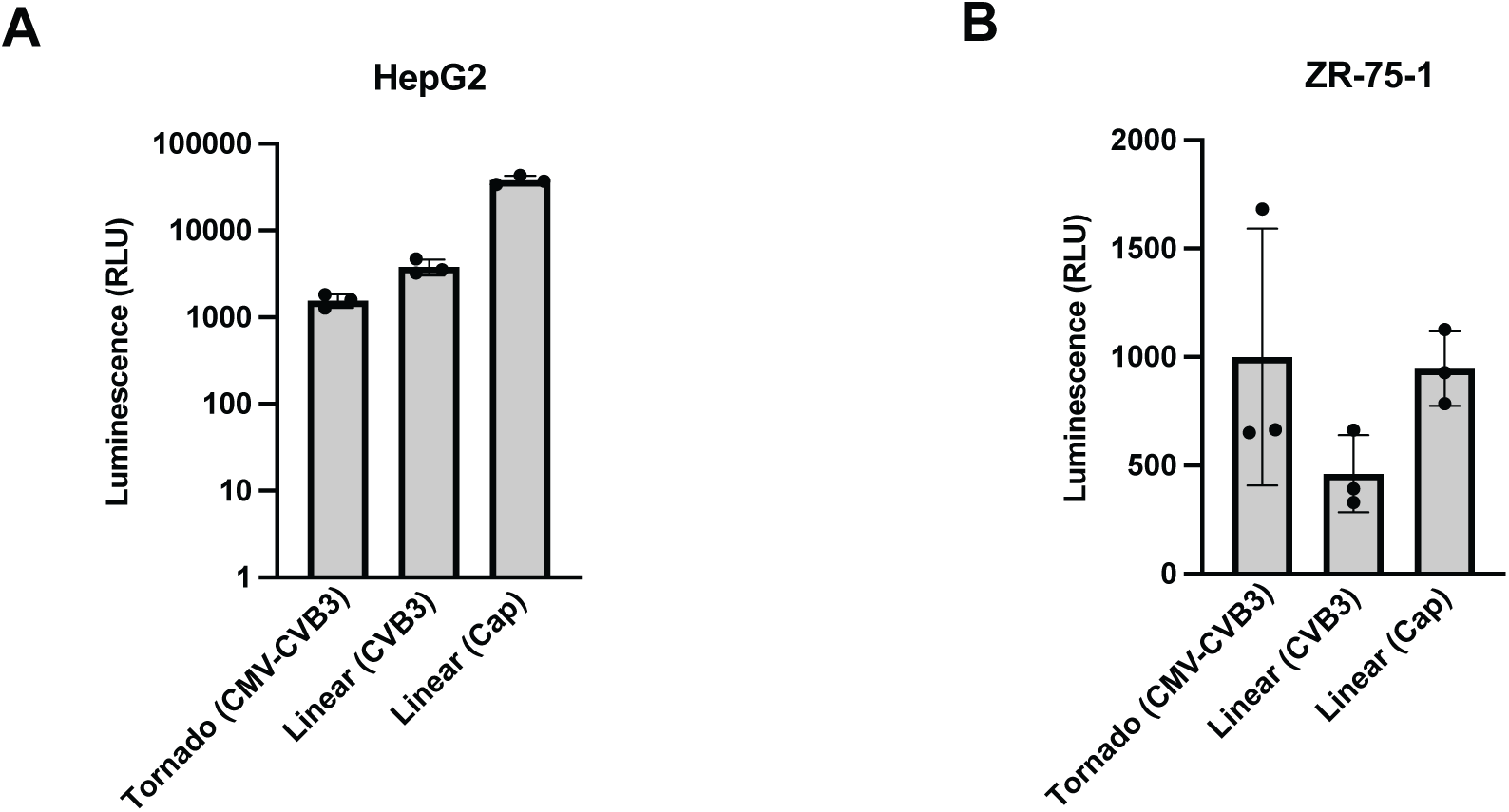
The Tornado translation system can be used in multiple cell types. (A), (B) The Tornado translation system can be used in multiple cell types. We quantified luminescence from HepG2 and ZR-75-1 cells transfected with plasmids expressing the Tornado translation system (Tornado (CMV-CVB3)), the linear cap-dependent mRNA expression system (Linear (Cap)), and the linear CVB3-dependent mRNA expression system (Linear (CVB3)). Tornado translation system produces luminescence in HepG2 (A) and ZR-75-1 (B) cells. Notably, the Tornado translation system produced similar levels of luminescence in ZR-75-1 cells as the linear cap-dependent mRNA expression system. RLU = Relative Luminescence units. Data are presented as mean values +/− one SD (n = 3 biological replicates). Significance was calculated using unpaired two-tailed student’s *t*-test. ****p<.0001, ***p<.001, **p<.01, *p<.05, n.s. p>.05.

**Figure S10.**
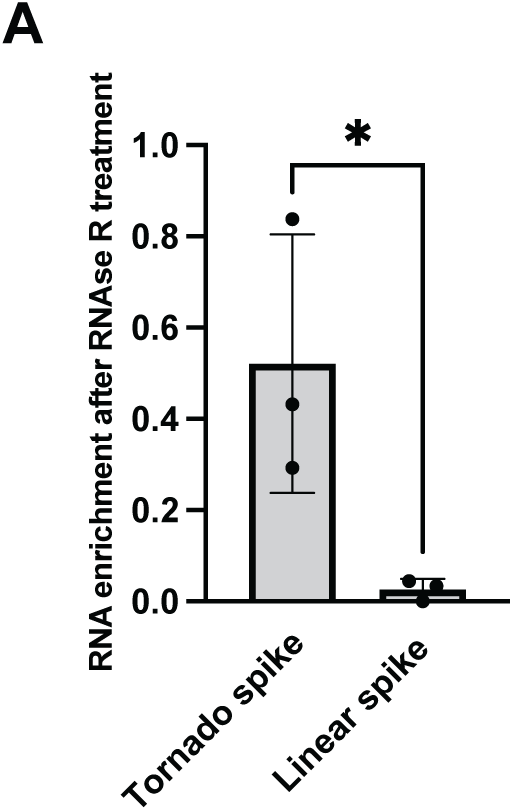
Expression of SARS-CoV-2 spike protein using the Tornado translation system. (A) The Tornado translation system can circularize the SARS-CoV-2 spike protein mRNA, which is longer that the nLuc mRNA. The total length of the SARS-CoV-2 mRNA with the CVB3 IRES is 4719 nt, compared to 1527 nt, which is the length of the split nLuc mRNA with the CVB3 IRES. HEK293T cells were transfected with plasmids expressing the Tornado translation system and the linear mRNA expression system containing a spike protein insert. RNA was treated with vehicle or RNase R then quantified by doing qRT-PCR with primers that amplified a 124 nt region of the spike protein. The Tornado translation system produces an RNA that is circular as evidenced by its resistance to RNase R compared to the linear control. Data are presented as mean values +/− one SD (n = 3 biological replicates). Significance was calculated using unpaired two-tailed student’s *t*-test. ****p<.0001, ***p<.001, **p<.01, *p<.05, n.s. p>.05.

**Figure S11, related to Figure 6.**
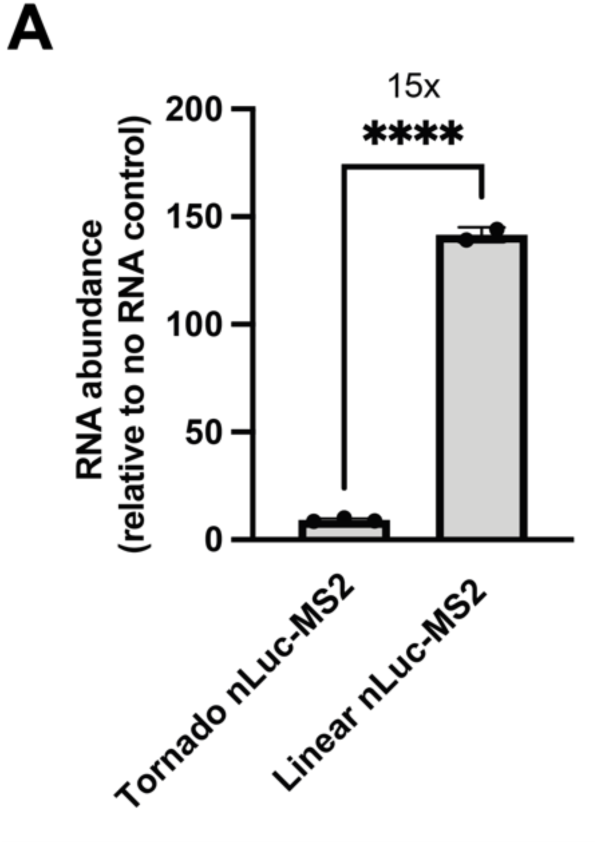
Titer of VLPs produced by using the Tornado translation system and the linear cap-dependent mRNA expression system. (A) VLP RNA titers. We titered the VLPs that were produced using the Tornado translation system and the linear mRNA expression system using RNA quantification. We extracted viral RNA from equal volumes of viral supernatant that was produced using the Tornado translation system (Tornado nLuc-MS2) and the linear mRNA expression system (Linear nLuc-MS2). We then performed qRT-PCR using primers that amplified a 126nt region of the nLuc gene. The linear mRNA expression system produced VLPs that were 15-fold more concentrated than the VLPs produced by the Tornado translation system. Data are presented as mean values +/− one SD (n = 3 biological replicates). Significance was calculated using unpaired two-tailed student’s *t*-test. ****p<.0001, ***p<.001, **p<.01, *p<.05, n.s. p>.05.

**Figure S12, related to Figure 6.**
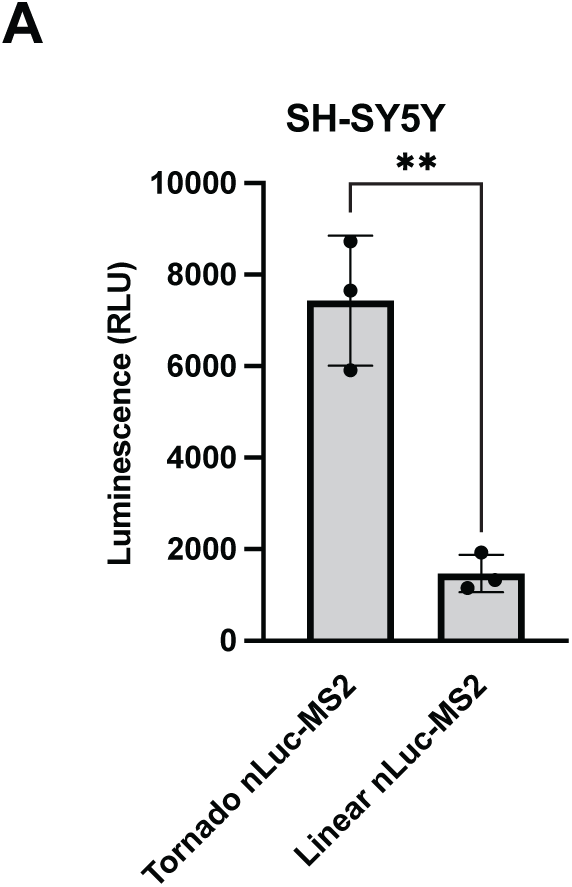
VLPs produced by using the Tornado translation system can be used on an alternative cell line. (A) Our previous experiments were performed in HEK293T cells. To ensure that other cells could be used, we transduced SH-SY5Y neuroblastoma cells with VLPs produced using the Tornado translation system. We quantified luminescence from SH-SY5Y cells transduced with VLPs that were produced using either the Tornado translation system (Tornado nLuc-MS2) or the linear mRNA expression system (Linear nLuc-MS2) at hour 24 after transduction. Cells were transduced with equal levels of VLP mRNA. VLPs that were produced using the Tornado translation system can be used to transduce SH-SY5Y cells and produce ∼5-fold more luminescence than the VLPs that were produced using the linear mRNA expression system. RLU = Relative Luminescence units. Data are presented as mean values +/− one SD (n = 3 biological replicates). Significance was calculated using unpaired two-tailed student’s *t*-test. ****p<.0001, ***p<.001, **p<.01, *p<.05, n.s. p>.05.

## REFERENCES

Abe, B.T., Wesselhoeft, R.A., Chen, R., Anderson, D.G., and Chang, H.Y. (2022). Circular RNA migration in agarose gel electrophoresis. Mol Cell 82, 1768–1777.e1763.

Abe, N., Hiroshima, M., Maruyama, H., Nakashima, Y., Nakano, Y., Matsuda, A., Sako, Y., Ito, Y., and Abe, H. (2013). Rolling circle amplification in a prokaryotic translation system using small circular RNA. Angew Chem Int Ed Engl 52, 7004–7008.

Abe, N., Matsumoto, K., Nishihara, M., Nakano, Y., Shibata, A., Maruyama, H., Shuto, S., Matsuda, A., Yoshida, M., Ito, Y., et al. (2015). Rolling Circle Translation of Circular RNA in Living Human Cells. Scientific Reports 5, 16435.

Bailey Jennifer, M., and Tapprich William, E. (2007). Structure of the 5′ Nontranslated Region of the Coxsackievirus B3 Genome:Chemical Modification and Comparative Sequence Analysis. Journal of Virology 81, 650–668.

Brooks, A.R., Harkins, R.N., Wang, P., Qian, H.S., Liu, P., and Rubanyi, G.M. (2004). Transcriptional silencing is associated with extensive methylation of the CMV promoter following adenoviral gene delivery to muscle. The Journal of Gene Medicine 6, 395–404.

Brown, E.A., Zhang, H., Ping, L.H., and Lemon, S.M. (1992). Secondary structure of the 5’ nontranslated regions of hepatitis C virus and pestivirus genomic RNAs. Nucleic Acids Res 20, 5041–5045.

Chen, C., Zhang, H., Broitman, S.L., Reiche, M., Farrell, I., Cooperman, B.S., and Goldman, Y.E. (2013). Dynamics of translation by single ribosomes through mRNA secondary structures. Nat Struct Mol Biol 20, 582–588.

Chen, C.-K., Cheng, R., Demeter, J., Chen, J., Weingarten-Gabbay, S., Jiang, L., Snyder, M.P., Weissman, J.S., Segal, E., Jackson, P.K., et al. (2021). Structured elements drive extensive circular RNA translation. Molecular Cell 81, 4300–4318.e4313.

Chen, C.-y., and Sarnow, P. (1995). Initiation of Protein Synthesis by the Eukaryotic Translational Apparatus on Circular RNAs. Science 268, 415–417.

Chen, R., Wang, S.K., Belk, J.A., Amaya, L., Li, Z., Cardenas, A., Abe, B.T., Chen, C.-K., Wender, P.A., and Chang, H.Y. (2022). Engineering circular RNA for enhanced protein production. Nature Biotechnology.

Chen, Z.Y., He, C.Y., Meuse, L., and Kay, M.A. (2004). Silencing of episomal transgene expression by plasmid bacterial DNA elements in vivo. Gene Therapy 11, 856–864.

Cocquerelle, C., Mascrez, B., Hétuin, D., and Bailleul, B. (1993). Mis-splicing yields circular RNA molecules. Faseb j 7, 155–160.

Costello, A., Lao, N.T., Barron, N., and Clynes, M. (2019). Continuous translation of circularized mRNA improves recombinant protein titer. Metab Eng 52, 284–292.

Cronin, J., Zhang, X.Y., and Reiser, J. (2005). Altering the tropism of lentiviral vectors through pseudotyping. Curr Gene Ther 5, 387–398.

Dieci, G., and Sentenac, A. (1996). Facilitated recycling pathway for RNA polymerase III. Cell 84, 245–252.

Dixon, A.S., Schwinn, M.K., Hall, M.P., Zimmerman, K., Otto, P., Lubben, T.H., Butler, B.L., Binkowski, B.F., Machleidt, T., Kirkland, T.A., et al. (2016). NanoLuc Complementation Reporter Optimized for Accurate Measurement of Protein Interactions in Cells. ACS Chemical Biology 11, 400–408.

Hamilton, J.R., Tsuchida, C.A., Nguyen, D.N., Shy, B.R., McGarrigle, E.R., Sandoval Espinoza, C.R., Carr, D., Blaeschke, F., Marson, A., and Doudna, J.A. (2021). Targeted delivery of CRISPR-Cas9 and transgenes enables complex immune cell engineering. Cell Rep 35, 109207.

Harger, J.W., and Dinman, J.D. (2003). An in vivo dual-luciferase assay system for studying translational recoding in the yeast Saccharomyces cerevisiae. Rna 9, 1019–1024.

Ho-Xuan, H., Glažar, P., Latini, C., Heizler, K., Haase, J., Hett, R., Anders, M., Weichmann, F., Bruckmann, A., Van den Berg, D., et al. (2020). Comprehensive analysis of translation from overexpressed circular RNAs reveals pervasive translation from linear transcripts. Nucleic Acids Research 48, 10368–10382.

Honda, M., Beard, M.R., Ping, L.H., and Lemon, S.M. (1999). A phylogenetically conserved stem-loop structure at the 5’ border of the internal ribosome entry site of hepatitis C virus is required for cap-independent viral translation. J Virol 73, 1165–1174.

Huang, Y., Yang, C., Xu, X.-f., Xu, W., and Liu, S.-w. (2020). Structural and functional properties of SARS-CoV-2 spike protein: potential antivirus drug development for COVID-19. Acta Pharmacologica Sinica 41, 1141–1149.

Ibrahim, H., Wilusz, J., and Wilusz, C.J. (2008). RNA recognition by 3’-to-5’ exonucleases: the substrate perspective. Biochim Biophys Acta 1779, 256–265.

Jang, S.K., and Wimmer, E. (1990). Cap-independent translation of encephalomyocarditis virus RNA: structural elements of the internal ribosomal entry site and involvement of a cellular 57-kD RNA-binding protein. Genes & Development 4, 1560–1572.

Jeck, W.R., Sorrentino, J.A., Wang, K., Slevin, M.K., Burd, C.E., Liu, J., Marzluff, W.F., and Sharpless, N.E. (2013). Circular RNAs are abundant, conserved, and associated with ALU repeats. Rna 19, 141–157.

Jiang, Y., Chen, X., and Zhang, W. (2021). Overexpression-based detection of translatable circular RNAs is vulnerable to coexistent linear RNA byproducts. Biochemical and Biophysical Research Communications 558, 189–195.

Kaminski, A.N.N., and Jackson, R.J. (1998). The polypyrimidine tract binding protein (PTB) requirement for internal initiation of translation of cardiovirus RNAs is conditional rather than absolute. RNA 4, 626–638.

Kozak, M. (2003). Alternative ways to think about mRNA sequences and proteins that appear to promote internal initiation of translation. Gene 318, 1–23.

Kozak, M. (2005). A second look at cellular mRNA sequences said to function as internal ribosome entry sites. Nucleic Acids Res 33, 6593–6602.

Liang, D., and Wilusz, J.E. (2014). Short intronic repeat sequences facilitate circular RNA production. Genes Dev 28, 2233–2247.

Litke, J.L., and Jaffrey, S.R. (2019). Highly efficient expression of circular RNA aptamers in cells using autocatalytic transcripts. Nature Biotechnology 37, 667–675.

Liu, Z., Chen, O., Wall, J.B.J., Zheng, M., Zhou, Y., Wang, L., Ruth Vaseghi, H., Qian, L., and Liu, J. (2017). Systematic comparison of 2A peptides for cloning multi-genes in a polycistronic vector. Scientific Reports 7, 2193.

Lu, B., Javidi-Parsijani, P., Makani, V., Mehraein-Ghomi, F., Sarhan, W.M., Sun, D., Yoo, K.W., Atala, Z.P., Lyu, P., and Atala, A. (2019). Delivering SaCas9 mRNA by lentivirus-like bionanoparticles for transient expression and efficient genome editing. Nucleic Acids Research 47, e44–e44.

Mäkinen, P.I., Koponen, J.K., Kärkkäinen, A.M., Malm, T.M., Pulkkinen, K.H., Koistinaho, J., Turunen, M.P., and Ylä-Herttuala, S. (2006). Stable RNA interference: comparison of U6 and H1 promoters in endothelial cells and in mouse brain. J Gene Med 8, 433–441.

Maugeri, M., Nawaz, M., Papadimitriou, A., Angerfors, A., Camponeschi, A., Na, M., Hölttä, M., Skantze, P., Johansson, S., Sundqvist, M., et al. (2019). Linkage between endosomal escape of LNP-mRNA and loading into EVs for transport to other cells. Nature Communications 10, 4333.

Naldini, L., Blömer, U., Gallay, P., Ory, D., Mulligan, R., Gage, F.H., Verma, I.M., and Trono, D. (1996). In Vivo Gene Delivery and Stable Transduction of Nondividing Cells by a Lentiviral Vector. Science 272, 263–267.

Obi, P., and Chen, Y.G. (2021). The design and synthesis of circular RNAs. Methods 196, 85–103.

Orioli, A., Pascali, C., Quartararo, J., Diebel, K.W., Praz, V., Romascano, D., Percudani, R., van Dyk, L.F., Hernandez, N., Teichmann, M., et al. (2011). Widespread occurrence of non-canonical transcription termination by human RNA polymerase III. Nucleic Acids Research 39, 5499–5512.

Pardi, N., Tuyishime, S., Muramatsu, H., Kariko, K., Mui, B.L., Tam, Y.K., Madden, T.D., Hope, M.J., and Weissman, D. (2015). Expression kinetics of nucleoside-modified mRNA delivered in lipid nanoparticles to mice by various routes. J Control Release 217, 345–351.

Prel, A., Caval, V., Gayon, R., Ravassard, P., Duthoit, C., Payen, E., Maouche-Chretien, L., Creneguy, A., Nguyen, T.H., Martin, N., et al. (2015). Highly efficient in vitro and in vivo delivery of functional RNAs using new versatile MS2-chimeric retrovirus-like particles. Molecular Therapy - Methods & Clinical Development 2, 15039.

Puttaraju, M., and Been, M.D. (1992). Group I permuted intron-exon (PIE) sequences self-splice to produce circular exons. Nucleic acids research 20, 5357–5364.

Qu, L., Yi, Z., Shen, Y., Lin, L., Chen, F., Xu, Y., Wu, Z., Tang, H., Zhang, X., Tian, F., et al. (2022). Circular RNA vaccines against SARS-CoV-2 and emerging variants. Cell 185, 1728–1744.e1716.

Segel, M., Lash, B., Song, J., Ladha, A., Liu, C.C., Jin, X., Mekhedov, S.L., Macrae, R.K., Koonin, E.V., and Zhang, F. (2021). Mammalian retrovirus-like protein PEG10 packages its own mRNA and can be pseudotyped for mRNA delivery. Science 373, 882–889.

Shah, P., Ding, Y., Niemczyk, M., Kudla, G., and Plotkin, Joshua B. (2013). Rate-Limiting Steps in Yeast Protein Translation. Cell 153, 1589–1601.

Sizova Daria, V., Kolupaeva Victoria, G., Pestova Tatyana, V., Shatsky Ivan, N., and Hellen Christopher, U.T. (1998). Specific Interaction of Eukaryotic Translation Initiation Factor 3 with the 5′ Nontranslated Regions of Hepatitis C Virus and Classical Swine Fever Virus RNAs. Journal of Virology 72, 4775-4782.

Stein, B.S., Gowda, S.D., Lifson, J.D., Penhallow, R.C., Bensch, K.G., and Engleman, E.G. (1987). pH-independent HIV entry into CD4-positive T cells via virus envelope fusion to the plasma membrane. Cell 49, 659–668.

Wen, J.-D., Lancaster, L., Hodges, C., Zeri, A.-C., Yoshimura, S.H., Noller, H.F., Bustamante, C., and Tinoco, I. (2008). Following translation by single ribosomes one codon at a time. Nature 452, 598–603.

Wesselhoeft, R.A., Kowalski, P.S., and Anderson, D.G. (2018). Engineering circular RNA for potent and stable translation in eukaryotic cells. Nature Communications 9, 2629.

Xue, Y., Zhou, Y., Wu, T., Zhu, T., Ji, X., Kwon, Y.S., Zhang, C., Yeo, G., Black, D.L., Sun, H., et al. (2009). Genome-wide analysis of PTB-RNA interactions reveals a strategy used by the general splicing repressor to modulate exon inclusion or skipping. Mol Cell 36, 996–1006.

Yang, D., Cheung, P., Sun, Y., Yuan, J., Zhang, H., Carthy, C.M., Anderson, D.R., Bohunek, L., Wilson, J.E., and McManus, B.M. (2003). A Shine-Dalgarno-like Sequence Mediates in Vitro Ribosomal Internal Entry and Subsequent Scanning for Translation Initiation of Coxsackievirus B3 RNA. Virology 305, 31–43.

Zheng, Q., Ryvkin, P., Li, F., Dragomir, I., Valladares, O., Yang, J., Cao, K., Wang, L.S., and Gregory, B.D. (2010). Genome-wide double-stranded RNA sequencing reveals the functional significance of base-paired RNAs in Arabidopsis. PLoS Genet 6, e1001141.

